# Placental cis-eQTL analysis identifies preeclampsia-dependent genetic regulation at promoters

**DOI:** 10.64898/2026.10.05.756656

**Authors:** Gamze Yazgeldi Gunaydin, Jaakko Leinonen, Sini Ezer, Eero Kajantie, Katja Kivinen, Seppo Heinonen, Anneli Pouta, FINNPEC Group, Hannele Laivuori, Satu Wedenoja, Juha Kere, Shintaro Katayama

**Author notes:** **Competing interests:** The authors declare no competing interests.

## Abstract

**Background:** Preeclampsia (PE) is a hypertensive pregnancy disorder with contributions from both maternal and fetal genetic factors, but how both genomes regulate placental transcription remains incompletely understood.

**Results:** We analyzed placental 5′-end RNA sequencing data from 59 pregnancies in the FINNPEC cohort and performed separate maternal and fetal cis-eQTL analyses using gene-level expression and promoter-level expression defined as Transcript Far 5’ Ends (TFE). Genotype-by-PE interaction models were used to identify genetic effects that differ according to PE status. No gene-level interaction eQTLs remained significant after multiple-testing correction, although suggestive signals included *LILRB2* in the maternal and *LPCAT3* in the fetal analyses. In contrast, we identified two significant syntenic promoter-level interaction eQTLs: rs12504642–TFE61820 at the *FHDC1* locus in the maternal analysis and rs1557150– TFE9032 at the *PPIAP31* locus in the fetal analysis.

**Conclusion:** These findings suggest that PE-dependent genetic interaction effects may be detectable at specific transcript promoter regions even when they are not captured at annotated gene levels.

## INTRODUCTION

Preeclampsia (PE) is a complex pregnancy-specific disorder affecting ∼2–5% of pregnancies and remains a leading cause of maternal and neonatal morbidity and mortality (Campbell et al., 2023; Jung et al., 2022). It is responsible for about 46,000 maternal and 500,000 fetal or newborn deaths globally each year (Cresswell et al., 2025). The placenta has a central role in its pathogenesis and placental dysfunction can lead to systemic inflammation and endothelial dysfunction in the mother. However, PE is heterogenous and results from interactions between placental, maternal, immunological and genetic factors.

Genetic studies have identified multiple loci associated with PE and hypertensive disorders of pregnancy, supporting a polygenic contribution to disease susceptibility. Both maternal and fetal genetic factors contribute to PE susceptibility, consistent with the complex interaction between maternal physiology and the fetal-derived placenta. Recent large-scale genome-wide association studies (GWAS) identified several loci related to blood pressure regulation, angiogenesis, trophoblast biology and immune function (Honigberg et al., 2023; Tyrmi et al., 2023). However, many disease-associated variants are located in non-coding transcribed regions, suggesting that their effects may involve gene regulation (Maurano et al., 2012). Expression quantitative trait locus (eQTL) analysis can link genetic variation to changes in gene expression and help interpret these functional effects of disease-associated variants (Albert & Kruglyak, 2015).

Placental eQTL studies have identified extensive genetic regulation linked to placental gene expression, including variants located in placenta-specific regulatory regions (Choi et al., 2024). The regulatory effects in placentas may also depend on context-dependent. For example, the fetal PE-associated variant rs4769613 near *FLT1* has been proposed to act as a conditional eQTL, with its association with placental *FLT1* expression differing between preeclamptic and normotensive pregnancies (Kikas et al., 2020, 2021; McGinnis et al., 2017).

Regulatory effects may also differ between specific transcriptional regions. Genetic effects on promoter usage can be highly context-specific and regulatory changes may not always be captured by measures of gene-level expression (Alasoo et al., 2019). Analysis of transcript far 5’end regions (TFEs), annotating active promoters, may therefore provide additional information on genetic regulation at specific transcription initiation regions, which may contribute to differential usage of transcript isoforms. A recent transcriptomic study has highlighted the extensive isoform diversity in the placenta, including previously unannotated transcripts and alternative first-exon usage (Bresnahan et al., 2026).

In this study, we investigated maternal and fetal genetic regulation of placental expression in normotensive and preeclamptic pregnancies from the FINNPEC cohort. Using 5’end focused Single-Cell Tagged Reverse Transcription (STRT) RNA sequencing, we analyzed both annotated gene-level 5’regions and de novo promoter regions. We performed separate maternal and fetal cis-eQTL analyses and tested genotype-by-PE interactions to identify genetic effects that differ according to PE status.

## MATERIAL AND METHODS

### Sampling and sample characteristics

Samples used in this study were obtained from the Finnish Genetics of Pre-eclampsia Consortium (FINNPEC) study, a Finnish case–control cohort recruited at five university hospitals. In the original cohort profile, FINNPEC included 1,450 women with PE and 1,065 control women without PE. Preeclampsia was defined as new-onset hypertension (systolic blood pressure ≥140 mmHg or diastolic ≥90 mmHg) and proteinuria after 20 weeks of gestation. Control samples were women without PE recruited from the same clinical settings. Placental tissue was collected after delivery from a subcohort of participants (Jääskeläinen et al., 2016). Placental biopsy samples were collected immediately after delivery from villous tissue beneath the basal plate, as previously described (Wedenoja et al., 2020).

For the present study, previously generated STRT RNA-seq data from 64 pregnancies with available placental samples were re-analyzed. Following quality control and removal of technical duplicates and outliers, 59 unique pregnancies were retained for the final analysis (Figure 1). Clinical characteristics of the participants are summarized in Table 1. Descriptive statistics were generated using the table1 package (v1.4.3)(Rich B., 2025) in R (v4.5.2)(R Core Team, 2025). Group comparisons were performed using Student’s t-test for normally distributed continuous variables, the Mann–Whitney U test for non-normally distributed continuous variables and the chi-square test for categorical variables. Normality was assessed using the Shapiro–Wilk test.

**Figure 1.**
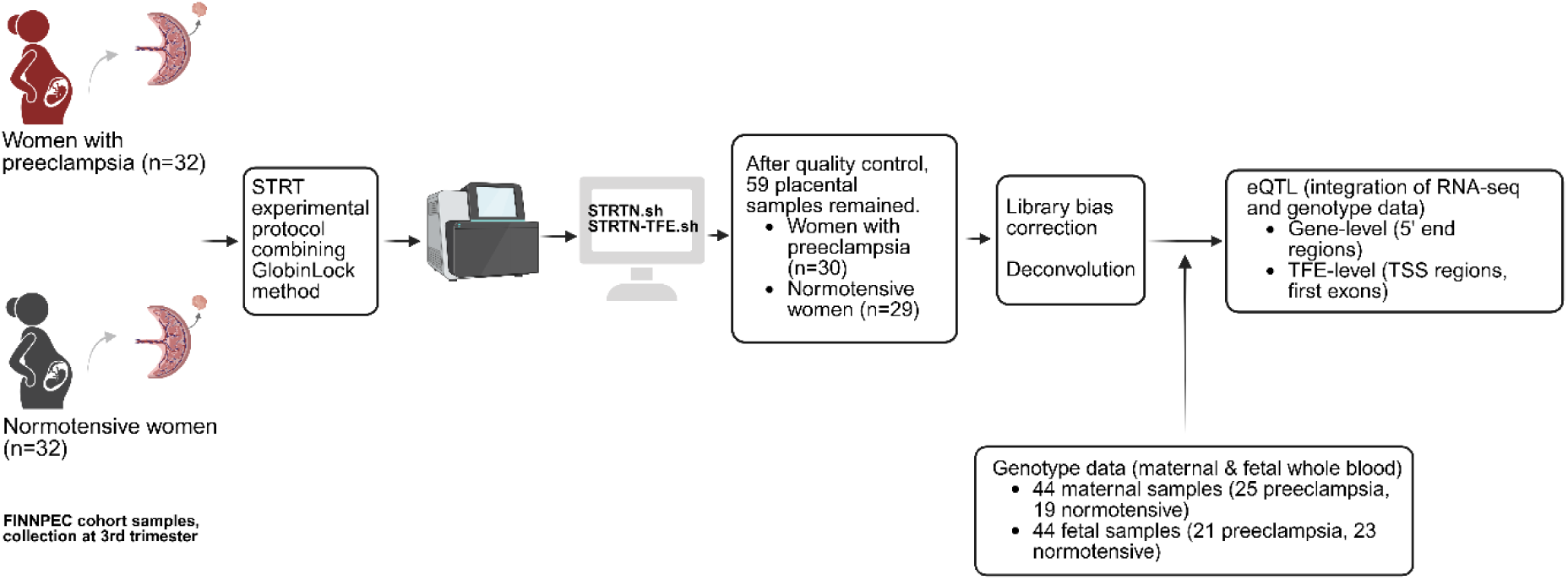
Overview of study. Study workflow showing the integration of placental STRT-seq expression profiles, maternal and fetal genotype dataset, clinical metadata and downstream analyses.

**Table 1.** Clinical characteristics of 59 pregnant women.

|  | NP<br>(N=29) | PE<br>(N=30) | P-value |
| --- | --- | --- | --- |
| Gestational week at birth |  |  |  |
| Mean (SD) | 38 (± 3.8) | 34 (± 4.9) | <0.001 |
| Premature birth |  |  |  |
| Term (>37) | 24 (83 %) | 11 (37 %) | <0.001 |
| Preterm (<37) | 5 (17 %) | 19 (63 %) |  |
| Delivery mode |  |  |  |
| Vaginal/Assisted Vaginal | 13 (45 %) | 8 (27 %) | <0.001 |
| Elective C-section | 11 (38 %) | 3 (10 %) |  |
| Semi-acute C-section | 5 (17 %) | 19 (63 %) |  |
| Child-sex |  |  |  |
| Boy | 19 (66 %) | 18 (60 %) | 0.866 |
| Girl | 10 (34 %) | 12 (40 %) |  |
| Birthweight standart deviation |  |  |  |
| Mean (SD) | -0.53 (± 1.3) | -1.2 (± 1.5) | 0.057 |
| Gravidity (number of pregnancies) |  |  |  |
| Mean (SD) | 1.9 (± 1.1) | 2.3 (± 3.0) | 0.58 |
| Parity (number of deliveries) |  |  |  |
| Mean (SD) | 0.62 (± 0.82) | 0.67 (± 2.2) | 0.152 |
| Preeclampsia history |  |  |  |
| No | 28 (97 %) | 25 (83 %) | 0.212 |
| Yes | 1 (3 %) | 5 (17 %) |  |
| Small for gestational age |  |  |  |
| No | 25 (86 %) | 18 (60 %) | 0.0488 |
| Yes | 4 (14 %) | 12 (40 %) |  |
| Body mass index |  |  |  |
| Mean (SD) | 26 (± 4.9) | 25 (± 4.7) | 0.331 |
| NA | 11 (37.9%) | 5 (16.7%) |  |
| Fetal HLA-G allele 1 |  |  |  |
| UTR1 | 15 (52 %) | 17 (57 %) | 0.663 |
| UTR2 | 10 (34 %) | 11 (37 %) |  |
| UTR4 | 4 (14 %) | 2 (7 %) |  |
| Fetal HLA-G allele 2 |  |  |  |
| UTR1 | 3 (10 %) | 6 (20 %) | 0.139 |
| UTR2 | 4 (14 %) | 7 (23 %) |  |
| UTR3 | 1 (3 %) | 4 (13 %) |  |
| UTR4 | 13 (45 %) | 9 (30 %) |  |
| UTR5 | 4 (14 %) | 0 (0 %) |  |
| UTR7 | 4 (14 %) | 2 (7 %) |  |
| UTR18 | 0 (0 %) | 1 (3 %) |  |
| UTR6 | 0 (0 %) | 1 (3 %) |  |
| Fetal HLA-G allele 1 evolutionary lineage 1, 2 or 3 |  |  |  |
| 1 | 23 (79 %) | 24 (80 %) | 1 |
| 2 | 6 (21 %) | 6 (20 %) |  |
| Fetal HLA-G allele 2 evolutionary lineage 1, 2 or 3 |  |  |  |
| 1 | 12 (41 %) | 12 (40 %) | 0.0911 |
| 2 | 13 (45 %) | 18 (60 %) |  |
| 3 | 4 (14 %) | 0 (0 %) |  |

### RNA extraction and STRT bulk RNA-seq library preparation

RNA extraction was performed as previously described (Wedenoja et al., 2020). The present study re-analyzed previously generated placental STRT bulk RNA-seq data from FINNPEC cohort. Two STRT RNA-seq libraries had been prepared as previously described (Wedenoja et al., 2020), using a modified 5′-end Single-cell Tagged Reverse Transcription (STRT) protocol combined with the GlobinLock technique (Krjutškov et al., 2016). A total of 40 ng RNA was used per sample, with 8-bp unique molecular identifiers (UMIs) and 6-bp sample-specific barcodes for multiplexing (Boskovic et al., 2023; Ezer et al., 2021). A total of 96 barcoded samples (48 per library) were generated from 64 placentas. The two libraries were sequenced on the Illumina NextSeq 500 platform using 75-cycle high-output runs with an 8M3S74T6B read structure. Samples were balanced for PE and NP status between the two libraries. ERCC RNA spike-in controls (97 transcripts) were included for normalization and quality control.

### STRT-seq data preprocessing and library bias correction

Raw sequencing data from the two STRT libraries were processed using the STRT pipeline (https://github.com/gyazgeldi/STRTN, commit e16a6d1), which generated gene-level and promoter-level expression matrices together with quality-control metrics, as previously described (Boskovic et al., 2023). Reads were aligned to the hg38 reference genome using GENCODE v43 basic annotation, which was downloaded from UCSC Genome Browser (https://genome.ucsc.edu/)(Frankish et al., 2021). Gene-level expression was quantified using annotated 5′ regions, including the 5′UTR and proximal upstream regions. Promoter-level expressions were quantified using transcription start site regions obtained from transcript assembly by StringTie (Pertea et al., 2015).

Sample quality was assessed using mapped reads, spike-in reads, spike-in 5′end rate, mapping rate, mapped/spike-in ratio and coding 5′end rate (Supplementary Table 1 and 2). Samples outside the Q1 − 1.5×IQR or Q3 + 1.5×IQR range were considered outliers. Twelve low-quality samples were excluded (Supplementary Figure 1), leaving 84 samples. Technical duplicates were then removed by retaining the sample with higher RNA quality, resulting in 59 unique placental samples for downstream analyses. Genes and promoters expressed in five or fewer samples were excluded from the analysis. Library bias between sequencing runs was corrected using a negative binomial generalized linear model (Supplementary Figure 2) (Katayama et al., 2019).

### Principal component analysis

Principal component analysis (PCA) was performed on library bias-corrected, spike-in-normalized and log-transformed expression data to assess major sources of variation. UMAP was used for two-dimensional visualization using Seurat (v5.4.0)(Hao et al., 2024). For downstream analyses, PCA was performed using PCAtools (v2.23.3)(Blighe K., Andrews J., Lun A., 2026) after removing the bottom 20% of low-variance features. Associations between expression PCs and clinical or technical variables were assessed to identify potential covariates for downstream analyses. Continuous variables were evaluated using Pearson correlation, whereas categorical variables were assessed using non-parametric tests. For visualization and correlation analyses, categorical variables were converted to binary indicators, including preeclampsia status, preterm birth, fetal sex, delivery mode and HLA-G 3′UTR diplotypes.

### Pathway enrichment analysis

Gene set enrichment analysis (GSEA) was performed using the curated pathway collection CP: KEGG_LEGACY (downloaded on 24 April 2026) (Kanehisa et al., 2025; Liberzon et al., 2015) to identify pathways associated with PE. Analyses were performed using the fgsea package (v1.36.2)(Korotkevich et al., 2016) in R. Genes were ranked based on principal component (PC) loading values, reflecting their contributions to each PC. Enrichment significance was assessed using normalized enrichment scores (NES) and adjusted p-values.

### Reference panel-based deconvolution

Cell-type deconvolution of bulk placental RNA-seq data was performed using MuSiC (v1.0.0) in R (Fan et al., 2022). Normalized bulk expression data were used as input, together with an external single-cell RNA-seq reference dataset GSE182381 derived from uncomplicated term villous placental tissue (≥37 weeks of gestation) and comprising 27 annotated cell types (Campbell et al., 2023). The reference dataset was converted to an object and cell-type proportions were estimated using the `music_prop` function. The estimated cell-type proportions were analyzed separately to characterize differences in placental cell-type composition between preeclampsia (PE) and normotensive (NP) samples and were not used to normalize the bulk expression data or included directly as covariates in the expression analyses. Cell types with zero variance or not represented in both groups were excluded.

Estimated cell-type proportions were compared between PE and NP groups using the Wilcoxon rank-sum test, followed by Benjamini–Hochberg correction for multiple testing. For visualization, estimated proportions were log10-transformed. Associations between estimated cell-type proportions, expression-derived PCs and clinical variables were also assessed using Pearson correlation for continuous variables and non-parametric tests for categorical variables and visualized using eigencorplot. For promoter-based analyses, annotated TFEs were grouped by gene and their counts were summed to generate gene-level expression profiles compatible with the single-cell reference panel. Unannotated TFEs were excluded from the deconvolution analysis.

### Genotyping and variant calling

Genotyping of the FINNPEC samples was performed using Infinium Global Screening Array-24 v2.0 BeadChip (Illumina Inc., San Diego, CA, USA) at Molecular Medicine Finland FIMM Technology Centre, University of Helsinki. Pre-imputation quality control of FINNPEC samples was carried out as described in Tyrmi et al. 2023 (Tyrmi et al., 2023), resulting in 2,521 mothers and 1,714 fetal samples with genome-wide SNP data available. The genotyped samples were imputed with Beagle v4.1 (Browning & Browning, 2016) using a population-specific reference panel SISu v3 based on 3,775 individuals of Finnish ancestry with whole genome sequencing data (Kurki et al., 2023).

### Expression quantitative trait loci analysis

Separate cis-eQTL interaction analyses were performed using maternal (n=44, 25 PE cases and 19 controls) and fetal (n=44, 21 PE cases and 23 controls) genotype datasets to investigate maternal and fetal genetic contributions to placental transcriptional regulation. Both gene-level expression based on annotated 5′ regions (12,546 genes) and promoter-level expression (67,360 TFEs) were analyzed using tensorQTL (Ongen et al., 2016; Taylor-Weiner et al., 2019).

Expression data were library bias-corrected, spike-in normalized and log10-transformed before analysis. Expression principal components (PCs) were evaluated for associations with clinical and technical variables including estimated placental cell-type proportions (Supplementary Figures 4 and 6). PCs associated with PE status were excluded from the covariate set to avoid removing disease-related biological variation. Estimated cell-type proportions were not directly included as covariates in the eQTL models. Expression PCs strongly correlated with genotype PCs were also excluded to reduce collinearity (Supplementary Figures 5 and 7). As a result of these exclusions, the gene-level maternal models included genotype PCs 1–10 and expression PCs 1, 3, 7, 9 and 10, whereas fetal models included genotype PCs 1–10 and expression PCs 2, 8, 9 and 10. Promoter-level maternal models included genotype PCs 1–10 and expression PCs 1, 4, 5, 7, 9 and 10, whereas fetal models included genotype PCs 1–10 and expression PCs 4, 5 and 6.

To identify PE-dependent genetic effects, genotype-by-PE interaction analyses were performed using tensorQTL with PE status as a binary interaction variable: `Expression ∼ genotype + PE status + genotype × PE status + covariates.` To correct for multiple testing in the gene-level and promoter-level analyses, the obtained nominal p-values for genotype-PE-interaction (pval_gi) were first corrected for the effective number of independent tests in each gene/locus, and these p-values were further adjusted using the Benjamini–Hochberg (BH) procedure to correct for the number of genes tested (N=12,546 in the gene-based and N=67,360 in the promoter-based analysis). The resulting BH adjusted p-values were used to determine statistical significance at a global threshold of false discovery rate (FDR) < 0.05.

### Functional interpretation

To functionally annotate candidate eQTL variants and promoter regions, we used the SCREEN database (Moore et al., 2026) to identify candidate cis-regulatory elements (cCREs), HaploReg (Ward et al., 2005) to examine regulatory annotations and the UCSC Genome Browser and FANTOM5 CAGE data to assess genomic context and transcription initiation signals. Identified eVariant–expression associations were also compared with GTEx eQTL data from other human tissues, as placental tissue is not represented in GTEx (GTEx Consortium, 2013).

## RESULTS

An overview of the study design and analytical workflow is shown in Figure 1. The demographic and clinical characteristics of the study population are summarized in Table 1.

After removal of technical duplicates and outlier samples, 59 placental samples were retained for downstream analyses. Sample characteristics are shown in Table 1. The observed statistically significant differences (p<0.05) between the NP and PE groups are consistent with impaired placental function. In line with this, lower birth weight and a higher frequency of small for gestational age (SGA) fetus were observed. Increased rates of preterm birth and cesarean section in the PE group likely reflect clinical management of these complications.

### Placental transcriptomic and cell-type alterations associated with PE

Reference-based deconvolution identified a lower estimated proportion of fetal Hofbauer cells in PE placentas, whereas no other cell-type differences remained significant after multiple-testing correction (Supplementary Figure 3). Several PE-associated expression PCs were also associated with estimated placental cell-type proportions (Supplementary Figures 4 and 6). PCA-GSEA of PE-associated gene expression PCs showed enrichment of immune-related pathways, including JAK–STAT signaling, antigen processing and presentation and leukocyte transendothelial migration (Supplementary Table 3).

### Gene-level interaction cis-eQTL analysis identifies suggestive PE-dependent regulatory signals

Gene-level cis-eQTL interaction analyses were performed separately for the maternal and fetal samples to identify genetic effects on placental expression that differed according to PE status. No gene-level genotype-by-PE interaction eQTLs remained significant after multiple-testing correction in both maternal and fetal analysis (Figure 2a, 2b). However, several suggestive interaction signals were observed.

**Figure 2.**
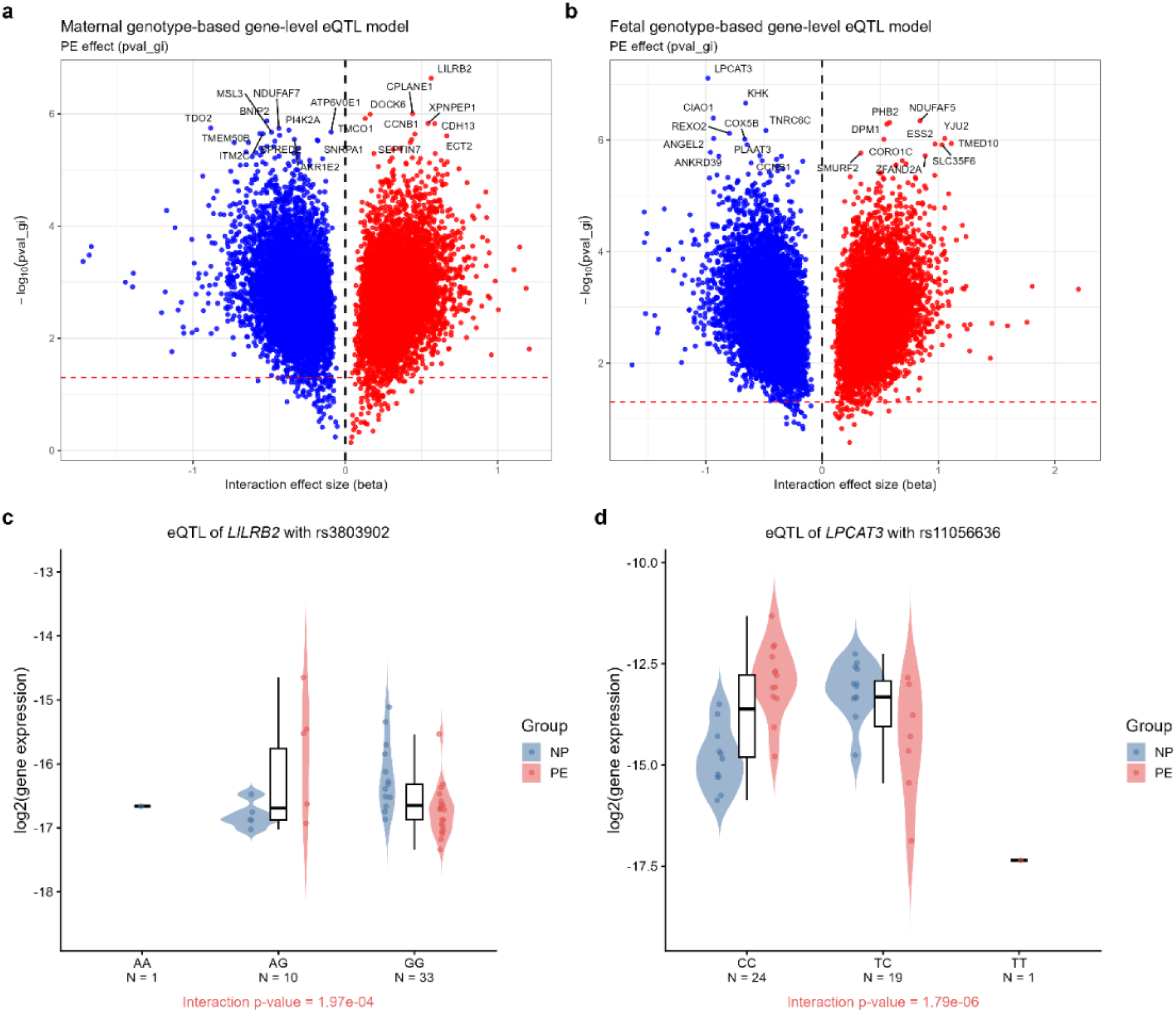
The strongest suggestive gene-level PE-dependent genetic effects were observed in for *LILRB2* (immune regulation), and *LPCAT3* (lipid metabolism). a, b) Volcano plots showing interaction eQTLs ranked by genotype × PE status interaction p-value (pval_gi) in the maternal and fetal genotype-based models, respectively. The x-axis shows the interaction effect size (b_gi), and the y-axis shows −log2 interaction pval_gi. Top 20 interaction eQTLs are labelled. c, d) Expression plots of the top interaction eQTLs from the maternal and fetal genotype-based models, showing *LILRB2* and *LPCAT3* expression, respectively. The x-axis shows genotype groups with sample numbers, and the y-axis shows spike-in normalized log2 gene expression. Samples are colored by PE status: PE in pink and NP in blue. The interaction p-value indicates whether the genotype effect on gene expression differs by PE status.

In the maternal analysis, the top variant based on the FDR-adjusted genotype-by-PE interaction p-value was rs35992881 near *MSL3* (pval_gi=2.1 × 10⁻⁶, FDR=0.21). A biologically relevant suggestive interaction signal was observed for rs3803902 linked to *LILRB2* (pval_gi=1.7 × 10⁻⁵, FDR=0.82), which encodes an immune receptor that binds MHC class I molecules, including HLA-G. Other top maternal signals included *CDH13, TDO2, DOCK6, CPLANE1* and *NDUFAF7* (all pval_gi<1.2 × 10⁻⁵, FD>0.81) (Figure 2a).

In the fetal analysis, rs35049337 near *KHK*, rs11681534 near *CIAO1* and rs11056636 near *LPCAT3* showed the lowest FDR values (pval_gi=2.2 × 10⁻⁷, 4.0 × 10⁻⁷ and 7.7 × 10⁻⁸, respectively; FDR = 0.14 for each). The rs11056636–*LPCAT3* interaction showed a negative interaction effect, consistent with a weaker genotype–expression association in PE than in NP pregnancies. Other top fetal signals included *PHB2, NDUFAF5, DPM1, ANGEL2, REXO* and *TNRC6C* (all pval_gi<4.3×10⁻⁶, FDR>0.14) (Figure 2b). Several top variants overlapped annotated regulatory features or showed eQTL evidence in other tissues (Supplementary Table 4).

### Promoter-level cis-eQTL interaction analysis identifies significant PE-dependent regulatory associations

After the gene-level analyses, promoter-level interaction analyses were performed separately for the maternal and fetal datasets. Two genotype-by-PE interaction eQTLs remained significant after multiple-testing correction. In the maternal analysis, rs12504642 was associated with TFE61820 at the *FHDC1* locus, with the lead variant located approximately 203 kb upstream of the TFE. In the fetal analysis, rs1557150 was associated with TFE9032 at the *PPIAP31* locus, with the lead variant located approximately 701 kb upstream of the TFE (Figure 3 and Table 2).

**Figure 3.**
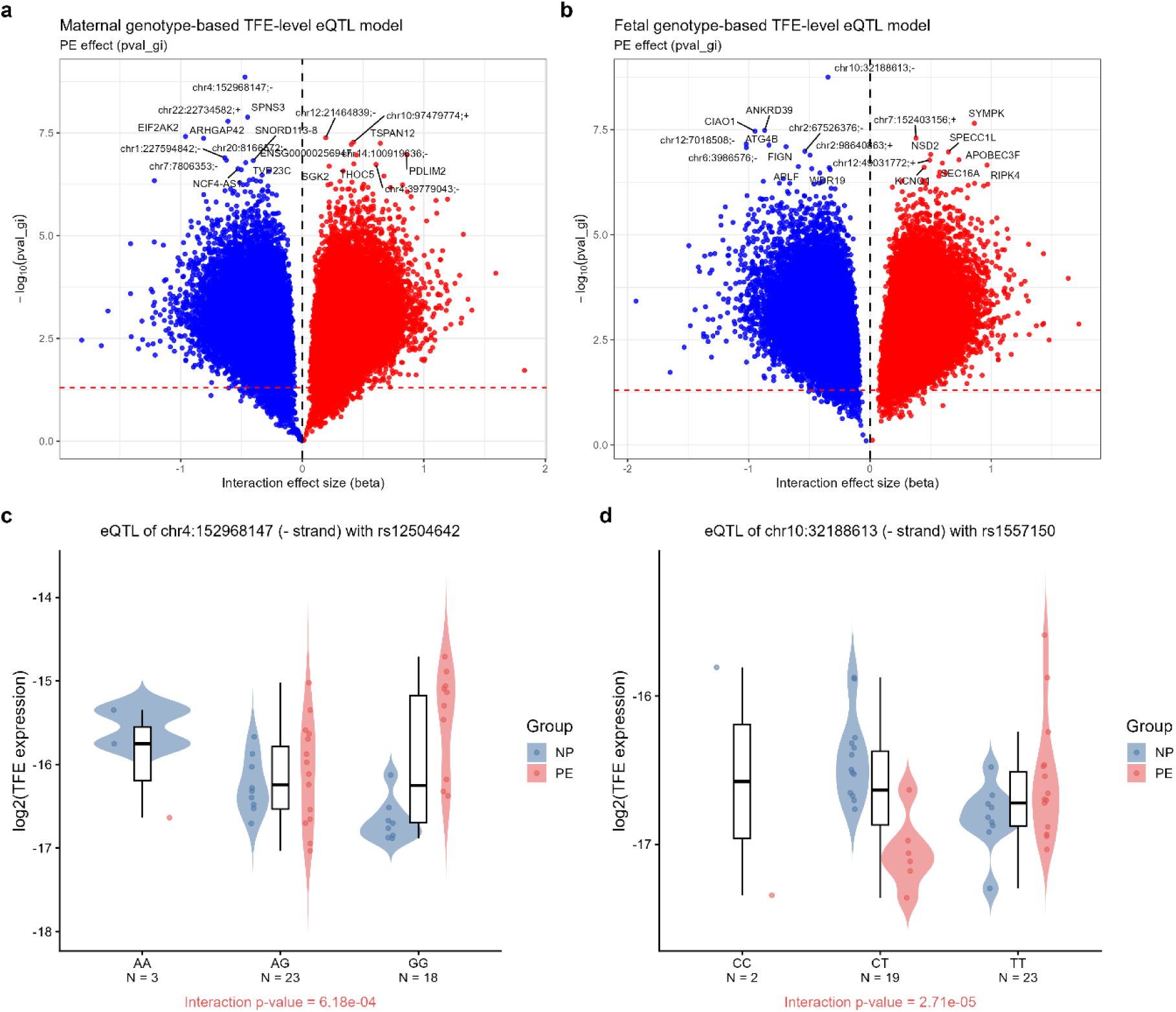
Promoter-level analysis reveals significant PE-dependent genetic regulation at specific transcriptional regions. a, b) Volcano plots showing interaction eQTLs ranked by genotype × PE status interaction p-value (pval_gi) in the maternal and fetal genotype-based models, respectively. The x-axis shows the interaction effect size (b_gi), and the y-axis shows −log10 interaction pval_gi. Top 20 interaction eQTLs are labelled. c, d) Expression plots of the top interaction eQTLs from the maternal and fetal genotype-based models, showing TFE61820 (chr4:152968147;-) and TFE9032 (chr10:32188613;-) expression, respectively. The x-axis shows genotype groups with sample numbers, and the y-axis shows spike-in normalized log10 gene expression. Samples are colored by PE status: PE in pink and NP in blue. The interaction p-value indicates whether the genotype effect on TFE expression differs by PE status.

**Table 2.**
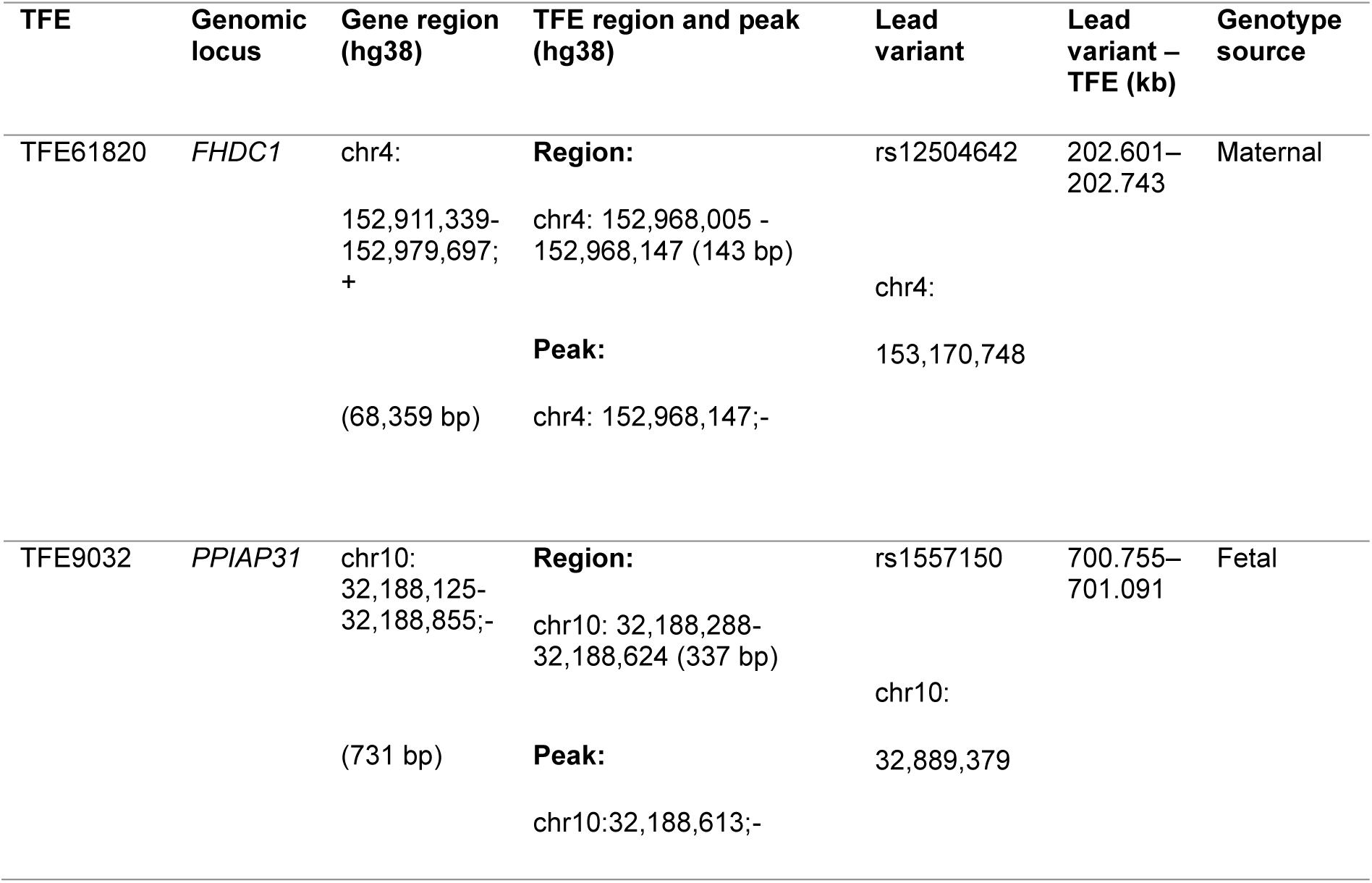
Genomic features of significant promoter-level interaction eQTLs.

Regulatory annotation provided additional genomic context for the two significant TFE interaction eQTLs. TFE61820 is located at chr4:152968005–152968147 on the negative strand, with a dominant 5′end peak at chr4:152968147. The maternal lead variant rs12504642 (chr4:153170748) is located approximately 202.6 kb from this peak. TFE61820 lies within an intronic region of *FHDC1*. The TFE peak overlaps the distal enhancer-like cCRE EH38E3610814 and predicted TFAP2A, TFAP2B, ZNF423 and ZNF454 binding motifs. A FANTOM5 CAGE-defined TSS was located 3 bp from the TFE peak and an additional CAGE signal was present within the TFE region. The lead maternal variant rs12504642 also overlapped a distal enhancer-like cCRE and an H3K27ac-enriched region. HaploReg further predicted allele-dependent changes in several transcription factor motifs at this locus (Figure 4).

**Figure 4.**
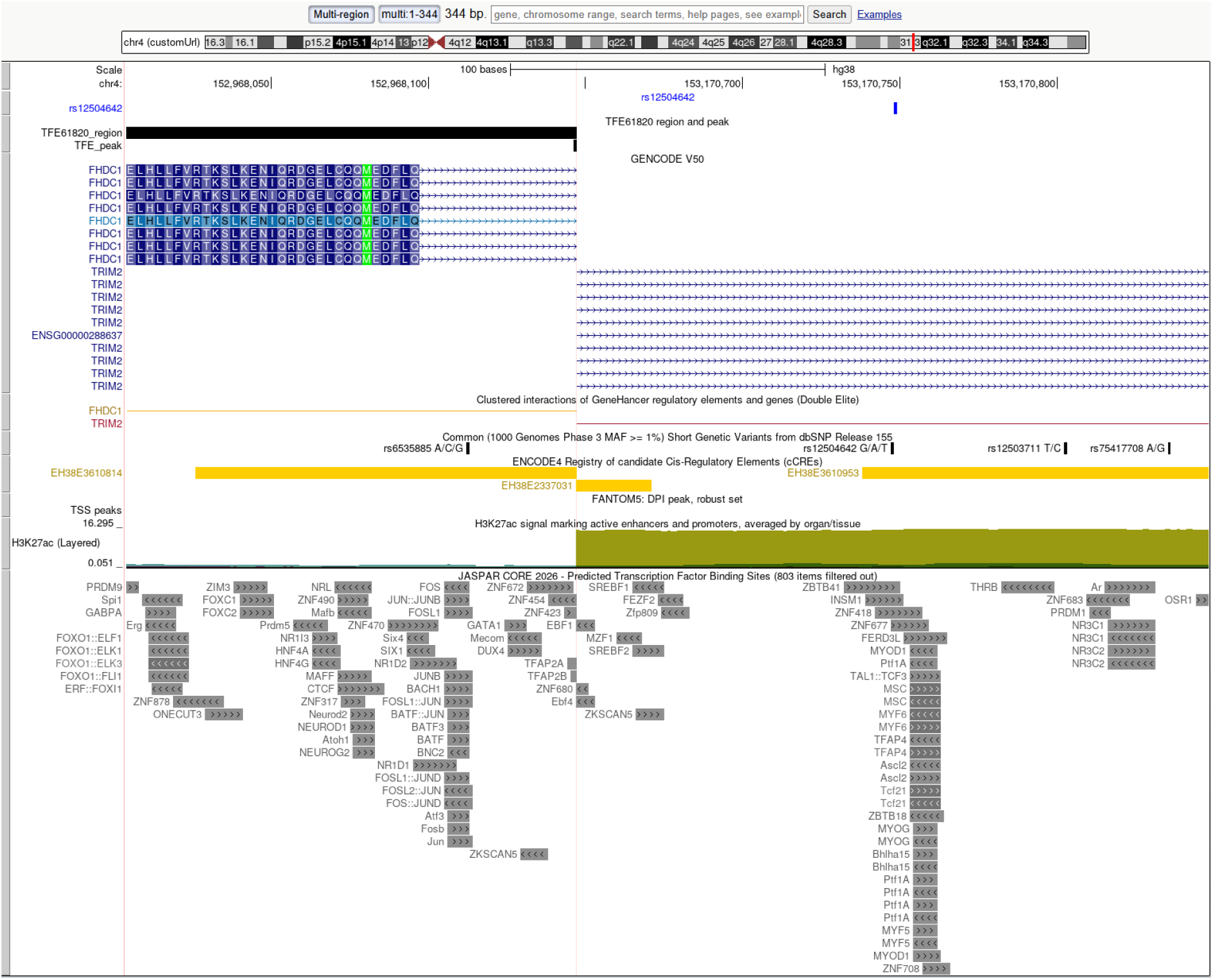
Genomic annotation of the detected maternal promoter interaction eQTL near *FHDC1*. The left panel shows TFE61820 (and its 1 bp TFE) peak located within the *FHDC1* locus, whereas the right panel shows the lead interaction SNP rs12504642 located within *TRIM2* locus and located approximately 203 kb upstream. UCSC Genome Browser tracks for GENCODE v50, GeneHancer regulatory element–gene interactions, dbSNP, ENCODE cCREs, FANTOM5 TSS peaks, H3K27ac and JASPAR predicted transcription factor binding motifs are shown.

Additionally, we also identified a significant interaction eQTL between rs1557150 and TFE9032. TFE9032 is located at chr10:32188288–32188624 on the negative strand, with a dominant peak at chr10:32188613. The fetal lead variant rs1557150 (chr10:32889379) is located approximately 700.8 kb from the TFE peak (Table 2). TFE9032 lies within the *PPIAP31* pseudogene region, whereas the lead interaction variant rs1557150 overlaps the ENCODE candidate cis-regulatory element EH38E4005881 (CA-CTCF class). The rs1557150 locus also overlaps an H3K27ac-enriched region. Both the TFE peak and the rs1557150 locus overlapped predicted JASPAR binding sites. However, several predicted motifs such as CREM, FOS::JUN and FOSL2::JUND were found near TFE9032 region (Figure 5). HaploReg v4.2 also predicted allele-specific changes in FOXA, IRF and STAT transcription factor motifs at rs1557150.

**Figure 5.**
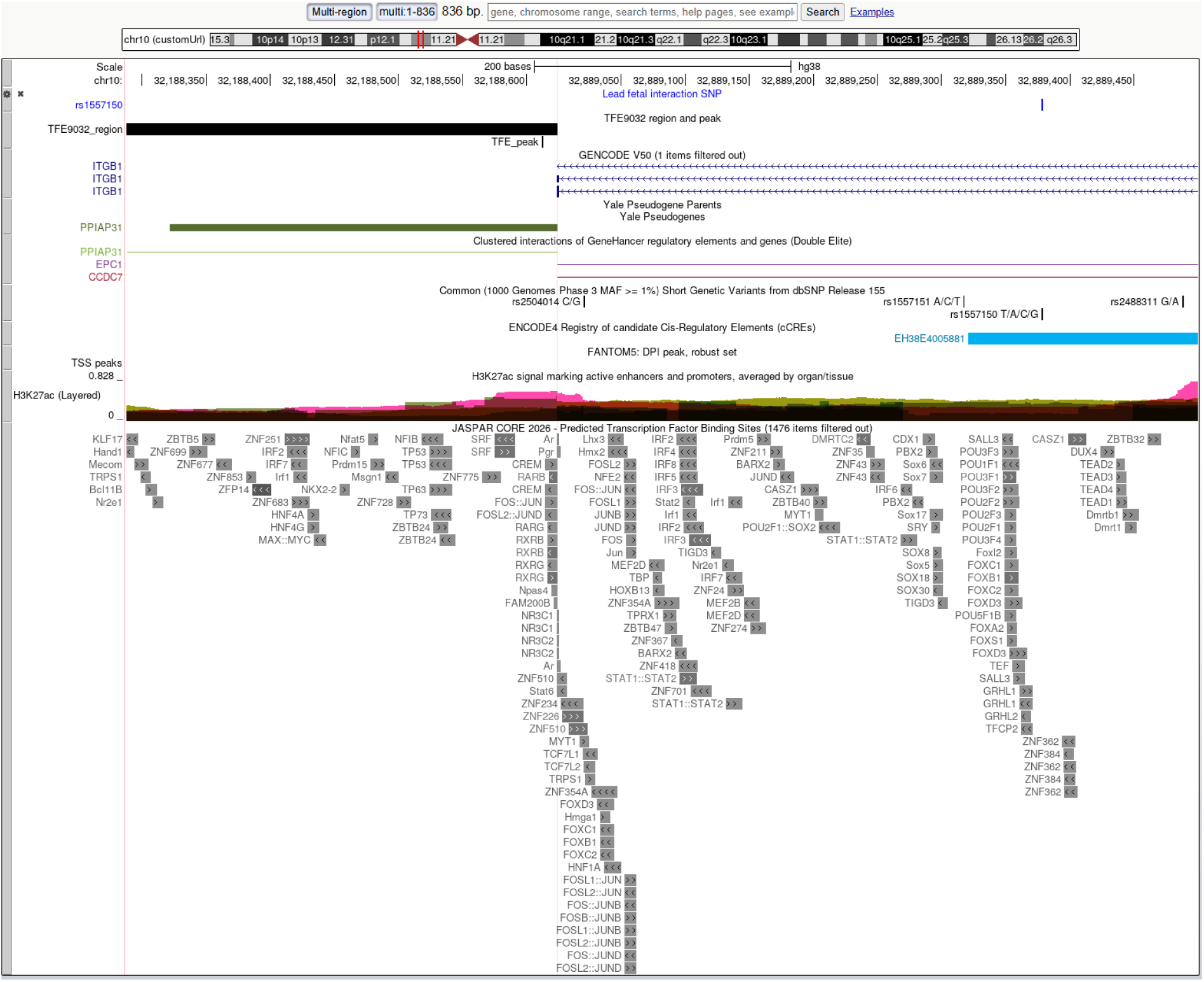
Genomic annotation of the significant fetal promoter interaction eQTL. This figure was generated using the UCSC Genome Browser multi-region mode. The left panel shows TFE9032 (including its 1 bp TFE peak), which overlaps the *PPIAP31* pseudogene, whereas the right panel shows the lead fetal interaction eQTL variant rs1557150, located approximately 701 kb upstream of TFE9032 peak. The figure includes GENCODE gene annotations, Yale Pseudogene annotations, GeneHancer regulatory element–gene interactions, ENCODE candidate cis-regulatory elements (cCREs), H3K27ac signal, FANTOM5 transcription start site (TSS) peaks, and JASPAR predicted transcription factor binding sites.

Together, these findings show that PE-dependent genetic effects can be detected at specific promoter regions even when significant interactions are not observed at annotated gene-level 5′ regions. Both statistically significant promoter-level loci showed regulatory features, supporting the possibility that genetic regulation differs between specific transcriptional regions according to PE status.

## DISCUSSION

Our findings suggest that the effects of maternal and fetal genetic variation on placental transcription may differ between normotensive and preeclamptic pregnancies. No gene-level genotype-by-PE interactions remained significant after multiple-testing correction, although several suggestive signals involved genes related to immune regulation and lipid metabolism. In contrast, promoter-level analysis identified two significant interaction eQTLs, indicating that PE-dependent genetic effects may be detectable at specific promoter regions even when they are not detected using annotated gene-level 5′ regions.

Among the suggestive gene-level interactions, the maternal signal involving *LILRB2* is of particular interest because *LILRB2* is an inhibitory receptor for HLA-G at the maternal–fetal interface (Shiroishi et al., 2006). The fetal analysis highlighted *LPCAT3*, which is involved in phospholipid remodeling and was also increased in preeclamptic placentas in our previous transcriptomic study (Kaartinen et al., 2025). However, because neither interaction remained significant after multiple-testing correction, these findings should be interpreted cautiously.

The maternal promoter interaction analysis identified TFE61820 within the *FHDC1* region. This TFE overlaps an enhancer-like candidate cis-regulatory element and lies close to a FANTOM5 CAGE-defined transcription start site, providing independent support for transcriptional activity at this region. Predicted binding motifs included TFAP2A and TFAP2B. TFAP2A has previously been linked to decidualization and regulation of *sFLT1* in preeclampsia and has recently been suggested among candidate transcription factors regulating placental sFLT1 expression (Manley et al., 2020; Yong et al., 2025). These annotations therefore support a possible regulatory function of the TFE61820 region, however further studies are needed.

The fetal promoter interaction analysis identified TFE9032 within the *PPIAP31* pseudogene region. *PPIAP31* is derived from *PPIA*, which encodes Cyclophilin A (CypA). Although the function of *PPIAP31* itself is unclear, CypA has been implicated in PE and experimentally shown to reduce trophoblast migration and invasion and induce PE-like features in mice (Hu et al., 2020). Unlike TFE61820, TFE9032 did not directly overlap a known cCRE or FANTOM5 CAGE signal in our annotation. The associated variant is located within an intronic region of *ITGB1*. *ITGB1* has an established role in trophoblast function, and reduced placental *ITGB1* expression has been reported in PE; experimental studies further show that *ITGB1* regulates trophoblast proliferation, migration and invasion (Li et al., 2026; Shu et al., 2021). However, it is unclear whether *ITGB1* is related to this interaction. The role of TFE9032 locus is also unclear. Therefore, further studies are needed.

The lower estimated proportion of fetal Hofbauer cells observed in PE placentas is consistent with previous reports of reduced Hofbauer cell abundance (Chen et al., 2026; Yu et al., 2025) and altered macrophage populations in preeclamptic placentas (Yang et al., 2017). Because estimated cell-type proportions were not directly included as covariates in the eQTL models, variation in placental cell-type composition may contribute to some of the observed genotype– expression associations.

A major limitation of this study is the relatively small sample size, which limits statistical power to detect eQTLs under the current settings. Therefore, the significant TFE associations reported here should also be interpreted with caution and replicated in larger placental cohorts, ideally combined with further functional validation. In addition, placental samples were collected at delivery and consisted of mixed cell populations. Since cell-type composition varied between PE and NP samples and was not directly adjusted for in the eQTL models, some of the detected signals may partly reflect differences in cellular composition. Despite these limitations, the detection of significant interactions at the promoter-level suggests that examining specific transcript 5′ end regions, or novel promoters, may provide additional information on PE-dependent regulatory effects that are not captured by annotated gene-level 5′ regions.

## DATA AVAILABILITY

Due to patient data restrictions, researchers interested in using the FINNPEC data must obtain approval from the FINNPEC steering committee. Data requests may be subject to further review by the Ethics Committee and may also be subject to individual participant consent.

## ACKNOWLEDGEMENTS

The authors wish to acknowledge CSC – IT Center for Science, Finland, for computational resources, and thank Auli Saarinen for skilled technical assistance.

## Notes

### Competing Interest Statement

The authors have declared no competing interest.

## REFERENCES

Alasoo, K., Rodrigues, J., Danesh, J., Freitag, D. F., Paul, D. S., & Gaffney, D. J. (2019). Genetic effects on promoter usage are highly context-specific and contribute to complex traits. eLife, 8, e41673. 10.7554/eLife.41673

Albert, F. W., & Kruglyak, L. (2015). The role of regulatory variation in complex traits and disease. Nature Reviews. Genetics, 16(4), 197–212. 10.1038/nrg3891

Blighe K., Andrews J., Lun A. (2026). PCAtools: PCAtools: Everything Principal Components Analysis (Version R package version 2.23.3) [Computer software]. https://github.com/kevinblighe/PCAtools

Boskovic, N., Yazgeldi, G., Ezer, S., Tervaniemi, M. H., Inzunza, J., Deligiannis, S. P., Yaşar, B., Skoog, T., Krjutškov, K., Katayama, S., & Kere, J. (2023). Optimized single-cell RNA sequencing protocol to study early genome activation in mammalian preimplantation development. STAR Protocols, 4(3), 102357. 10.1016/j.xpro.2023.102357

Bresnahan, S. T., Yong, H. E. J., Nemani, A., Wu, W. H., Lopez, S., Chan, J. K. Y., White, F., Jacques, P.-É., Hivert, M.-F., Chan, S.-Y., Love, M. I., Huang, J. Y., & Bhattacharya, A. (2026). Long-read assembly reveals vast transcriptional complexity in the placenta associated with metabolic and endocrine function. Nature Communications, 17(1), 4795. 10.1038/s41467-026-71303-4

Browning, B. L., & Browning, S. R. (2016). Genotype Imputation with Millions of Reference Samples. The American Journal of Human Genetics, 98(1), 116–126. 10.1016/j.ajhg.2015.11.020

Campbell, K. A., Colacino, J. A., Puttabyatappa, M., Dou, J. F., Elkin, E. R., Hammoud, S. S., Domino, S. E., Dolinoy, D. C., Goodrich, J. M., Loch-Caruso, R., Padmanabhan, V., & Bakulski, K. M. (2023). Placental cell type deconvolution reveals that cell proportions drive preeclampsia gene expression differences. Communications Biology, 6(1), 264. 10.1038/s42003-023-04623-6

Chen, Y., Wu, L., Huang, D., Wu, X., Liu, K., Sun, B., Yang, J., Zhang, B., Ouyang, Z., Zhang, C., Tan, L., & Niu, J. (2026). Subtype specific immune-metabolic reprogramming in preeclampsia revealed by multiomics and serum biomarkers. Hypertension Research, 49(3), 641–657. 10.1038/s41440-025-02504-5

Choi, J., Lee, S. M., Norwitz, E. R., Kim, J. H., Jung, Y. M., Park, C.-W., Jun, J. K., Lee, D., Jin, Y., Kim, S., Cha, B., Park, J. S., & Kim, J.-I. (2024). Placental expression quantitative trait loci in an East Asian population. HGG Advances, 5(2), 100276. 10.1016/j.xhgg.2024.100276

Cresswell, J. A., Alexander, M., Chong, M. Y. C., Link, H. M., Pejchinovska, M., Gazeley, U., Ahmed, S. M. A., Chou, D., Moller, A.-B., Simpson, D., Alkema, L., Villanueva, G., Sguassero, Y., Tunçalp, Ö., Long, Q., Xiao, S., & Say, L. (2025). Global and regional causes of maternal deaths 2009-20: A WHO systematic analysis. The Lancet. Global Health, 13(4), e626–e634. 10.1016/S2214-109X(24)00560-6

Ezer, S., Yoshihara, M., Katayama, S., DoGA consortium, Daub, C., Lohi, H., Krjutskov, K., & Kere, J. (2021). Generation of RNA sequencing libraries for transcriptome analysis of globin-rich tissues of the domestic dog. STAR Protocols, 2(4), 100995. 10.1016/j.xpro.2021.100995

Fan, J., Lyu, Y., Zhang, Q., Wang, X., Li, M., & Xiao, R. (2022). MuSiC2: Cell-type deconvolution for multi-condition bulk RNA-seq data. Briefings in Bioinformatics, 23(6), bbac430. 10.1093/bib/bbac430

Frankish, A., Diekhans, M., Jungreis, I., Lagarde, J., Loveland, J. E., Mudge, J. M., Sisu, C., Wright, J. C., Armstrong, J., Barnes, I., Berry, A., Bignell, A., Boix, C., Carbonell Sala, S., Cunningham, F., Di Domenico, T., Donaldson, S., Fiddes, I. T., García Girón, C., … Flicek, P. (2021). GENCODE 2021. Nucleic Acids Research, 49(D1), D916–D923. 10.1093/nar/gkaa1087

GTEx Consortium. (2013). The Genotype-Tissue Expression (GTEx) project. Nature Genetics, 45(6), 580–585. 10.1038/ng.2653

Hao, Y., Stuart, T., Kowalski, M. H., Choudhary, S., Hoffman, P., Hartman, A., Srivastava, A., Molla, G., Madad, S., Fernandez-Granda, C., & Satija, R. (2024). Dictionary learning for integrative, multimodal and scalable single-cell analysis. Nature Biotechnology, 42(2), 293–304. 10.1038/s41587-023-01767-y

Honigberg, M. C., Truong, B., Khan, R. R., Xiao, B., Bhatta, L., Vy, H. M. T., Guerrero, R. F., Schuermans, A., Selvaraj, M. S., Patel, A. P., Koyama, S., Cho, S. M. J., Vellarikkal, S. K., Trinder, M., Urbut, S. M., Gray, K. J., Brumpton, B. M., Patil, S., Zöllner, S., … Natarajan, P. (2023). Polygenic prediction of preeclampsia and gestational hypertension. Nature Medicine, 29(6), 1540–1549. 10.1038/s41591-023-02374-9

Hu, H., Jiang, J., Chen, Q., Wei, S., Liu, M., Chen, X., Fan, C., Ma, J., Chen, W., Wang, X., & Zhong, M. (2020). Cyclophilin A inhibits trophoblast migration and invasion in vitro and vivo through p38/ERK/JNK pathways and causes features of preeclampsia in mice. Life Sciences, 261, 118351. 10.1016/j.lfs.2020.118351

Jääskeläinen, T., Heinonen, S., Kajantie, E., Kere, J., Kivinen, K., Pouta, A., Laivuori, H., & FINNPEC Study Group. (2016). Cohort profile: The Finnish Genetics of Pre-eclampsia Consortium (FINNPEC). BMJ Open, 6(11), e013148. 10.1136/bmjopen-2016-013148

Jung, E., Romero, R., Yeo, L., Gomez-Lopez, N., Chaemsaithong, P., Jaovisidha, A., Gotsch, F., & Erez, O. (2022). The etiology of preeclampsia. American Journal of Obstetrics and Gynecology, 226(2S), S844–S866. 10.1016/j.ajog.2021.11.1356

Kaartinen, L., Jääskeläinen, T., Sliz, E., Yazgeldi Gunaydin, G., Wedenoja, S., Katayama, S., Kajantie, E., Rinne, V., Heinonen, S., Kere, J., Merikallio, H., Hannele Laivuori submitted on behalf of FINNPEC group, Sliz, E., submitted on behalf of FinnGen group, Laivuori, H., & Hukkanen, J. (2025). Role of oxysterol 4β-hydroxycholesterol and liver X receptor alleles in pre-eclampsia. Annals of Medicine, 57(1), 2495763. 10.1080/07853890.2025.2495763

Kanehisa, M., Furumichi, M., Sato, Y., Matsuura, Y., & Ishiguro-Watanabe, M. (2025). KEGG: Biological systems database as a model of the real world. Nucleic Acids Research, 53(D1), D672–D677. 10.1093/nar/gkae909

Katayama, S., Skoog, T., Söderhäll, C., Einarsdottir, E., Krjutškov, K., & Kere, J. (2019). Guide for library design and bias correction for large-scale transcriptome studies using highly multiplexed RNAseq methods. BMC Bioinformatics, 20(1), 418. 10.1186/s12859-019-3017-9

Kikas, T., Inno, R., Ratnik, K., Rull, K., & Laan, M. (2020). C-allele of rs4769613 Near FLT1 Represents a High-Confidence Placental Risk Factor for Preeclampsia. Hypertension, 76(3), 884–891. 10.1161/HYPERTENSIONAHA.120.15346

Kikas, T., Laan, M., & Kasak, L. (2021). Current knowledge on genetic variants shaping placental transcriptome and their link to gestational and postnatal health. Placenta, 116, 2–11. 10.1016/j.placenta.2021.02.009

Korotkevich, G., Sukhov, V., Budin, N., Shpak, B., Artyomov, M. N., & Sergushichev, A. (2016). Fast gene set enrichment analysis. Bioinformatics. 10.1101/060012

Krjutškov, K., Koel, M., Roost, A. M., Katayama, S., Einarsdottir, E., Jouhilahti, E.-M., Söderhäll, C., Jaakma, Ü., Plaas, M., Vesterlund, L., Lohi, H., Salumets, A., & Kere, J. (2016). Globin mRNA reduction for whole-blood transcriptome sequencing. Scientific Reports, 6, 31584. 10.1038/srep31584

Kurki, M. I., Karjalainen, J., Palta, P., Sipilä, T. P., Kristiansson, K., Donner, K. M., Reeve, M. P., Laivuori, H., Aavikko, M., Kaunisto, M. A., Loukola, A., Lahtela, E., Mattsson, H., Laiho, P., Della Briotta Parolo, P., Lehisto, A. A., Kanai, M., Mars, N., Rämö, J., … Palotie, A. (2023). FinnGen provides genetic insights from a well-phenotyped isolated population. Nature, 613(7944), 508–518. 10.1038/s41586-022-05473-8

Li, X.-J., Wang, N., Jiang, F., Yang, J.-C., Wang, Y.-Y., & Wang, K. (2026). UBE3A Inhibits Trophoblast Cell Migration and Invasion by Promoting ITGB1 Degradation and Affecting PI3K/AKT Signaling. The Kaohsiung Journal of Medical Sciences, 42(4), e70122. 10.1002/kjm2.70122

Liberzon, A., Birger, C., Thorvaldsdóttir, H., Ghandi, M., Mesirov, J. P., & Tamayo, P. (2015). The Molecular Signatures Database (MSigDB) hallmark gene set collection. Cell Systems, 1(6), 417–425. 10.1016/j.cels.2015.12.004

Manley, C. N., Deepak, V., Ravikumar, N., Smith, A. K., Knight, A. K., Badell, M. L., Sidell, N., & Rajakumar, A. (2020). Transcription factor AP2A affects sFLT1 expression and decidualization in decidual stromal cells: Implications to preeclampsia pathology. Pregnancy Hypertension, 21, 152–158. 10.1016/j.preghy.2020.06.001

Maurano, M. T., Humbert, R., Rynes, E., Thurman, R. E., Haugen, E., Wang, H., Reynolds, A. P., Sandstrom, R., Qu, H., Brody, J., Shafer, A., Neri, F., Lee, K., Kutyavin, T., Stehling-Sun, S., Johnson, A. K., Canfield, T. K., Giste, E., Diegel, M., … Stamatoyannopoulos, J. A. (2012). Systematic localization of common disease-associated variation in regulatory DNA. Science, 337(6099), 1190–1195. 10.1126/science.1222794

McGinnis, R., Steinthorsdottir, V., Williams, N. O., Thorleifsson, G., Shooter, S., Hjartardottir, S., Bumpstead, S., Stefansdottir, L., Hildyard, L., Sigurdsson, J. K., Kemp, J. P., Silva, G. B., Thomsen, L. C. V., Jääskeläinen, T., Kajantie, E., Chappell, S., Kalsheker, N., Moffett, A., Hiby, S., … Morgan, L. (2017). Variants in the fetal genome near FLT1 are associated with risk of preeclampsia. Nature Genetics, 49(8), 1255–1260. 10.1038/ng.3895

Moore, J. E., Pratt, H. E., Fan, K., Phalke, N., Fisher, J., Elhajjajy, S. I., Andrews, G., Gao, M., Shedd, N., Fu, Y., Lacadie, M. C., Meza, J., Khandpekar, M., Ganna, M., Choudhury, E., Swofford, R., Phan, H., Ramirez, C. C., Campbell, M., … Weng, Z. (2026). An expanded registry of candidate cis-regulatory elements. Nature, 1–10. 10.1038/s41586-025-09909-9

Ongen, H., Buil, A., Brown, A. A., Dermitzakis, E. T., & Delaneau, O. (2016). Fast and efficient QTL mapper for thousands of molecular phenotypes. Bioinformatics, 32(10), 1479–1485. 10.1093/bioinformatics/btv722

Pertea, M., Pertea, G. M., Antonescu, C. M., Chang, T.-C., Mendell, J. T., & Salzberg, S. L. (2015). StringTie enables improved reconstruction of a transcriptome from RNA-seq reads. Nature Biotechnology, 33(3), 290–295. 10.1038/nbt.3122

R Core Team. (2025). R: A Language and Environment for Statistical Computing (Version 4.5.2) [R]. R Foundation for Statistical Computing. https://www.R-project.org/

Rich B. (2025). table1: Tables of Descriptive Statistics in HTML (Version R package version 1.5.1) [Computer software]. 10.32614/CRAN.package.table1

Shiroishi, M., Kuroki, K., Rasubala, L., Tsumoto, K., Kumagai, I., Kurimoto, E., Kato, K., Kohda, D., & Maenaka, K. (2006). Structural basis for recognition of the nonclassical MHC molecule HLA-G by the leukocyte Ig-like receptor B2 (LILRB2/LIR2/ILT4/CD85d). Proceedings of the National Academy of Sciences of the United States of America, 103(44), 16412–16417. 10.1073/pnas.0605228103

Shu, C., Han, S., Hu, C., Chen, C., Qu, B., He, J., Dong, S., & Xu, P. (2021). Integrin β1 regulates proliferation, apoptosis, and migration of trophoblasts through activation of phosphoinositide 3 kinase/protein kinase B signaling. The Journal of Obstetrics and Gynaecology Research, 47(7), 2406–2416. 10.1111/jog.14782

Taylor-Weiner, A., Aguet, F., Haradhvala, N. J., Gosai, S., Anand, S., Kim, J., Ardlie, K., Van Allen, E. M., & Getz, G. (2019). Scaling computational genomics to millions of individuals with GPUs. Genome Biology, 20(1), 228. 10.1186/s13059-019-1836-7

Tyrmi, J. S., Kaartokallio, T., Lokki, A. I., Jääskeläinen, T., Kortelainen, E., Ruotsalainen, S., Karjalainen, J., Ripatti, S., Kivioja, A., Laisk, T., Kettunen, J., Pouta, A., Kivinen, K., Kajantie, E., Heinonen, S., Kere, J., Laivuori, H., & FINNPEC Study Group, FinnGen Project, and the Estonian Biobank Research Team. (2023). Genetic Risk Factors Associated With Preeclampsia and Hypertensive Disorders of Pregnancy. JAMA Cardiology, 8(7), 674–683. 10.1001/jamacardio.2023.1312

Ward, K. N., Thiruchelvam, A. D., & Couto-Parada, X. (2005). Unexpected occasional persistence of high levels of HHV-6 DNA in sera: Detection of variants A and B. Journal of Medical Virology, 76(4), 563–570. 10.1002/jmv.20399

Wedenoja, S., Yoshihara, M., Teder, H., Sariola, H., Gissler, M., Katayama, S., Wedenoja, J., Häkkinen, I. M., Ezer, S., Linder, N., Lundin, J., Skoog, T., Sahlin, E., Iwarsson, E., Pettersson, K., Kajantie, E., Mokkonen, M., Heinonen, S., Laivuori, H., … Kere, J. (2020). Fetal HLA-G mediated immune tolerance and interferon response in preeclampsia. EBioMedicine, 59, 102872. 10.1016/j.ebiom.2020.102872

Yang, S. W., Cho, E. H., Choi, S. Y., Lee, Y. K., Park, J. H., Kim, M. K., Park, J. Y., Choi, H. J., Lee, J. I., Ko, H. M., Park, S. H., Hwang, H. S., & Kang, Y. S. (2017). DC-SIGN expression in Hofbauer cells may play an important role in immune tolerance in fetal chorionic villi during the development of preeclampsia. Journal of Reproductive Immunology, 124, 30–37. 10.1016/j.jri.2017.09.012

Yong, Q., van der Keur, C., Anholts, J. D. H., Kapsenberg, H., Mei, H., Bruijn, J. A., Eikmans, M., & Baelde, H. J. (2025). Identification of transcription factors that regulate placental sFLT1 expression. Molecular Human Reproduction, 31(3), gaaf031. 10.1093/molehr/gaaf031

Yu, X., Xiang, H., & Huang, X. (2025). Single-Cell Transcriptome and RNA Sequencing Reveal Immune-Related Markers of Preeclampsia. Reproductive Sciences, 32(8), 2819–2828. 10.1007/s43032-025-01843-5

