## Supplementary Figures and Tables for "Placental cis-eQTL analysis identifies preeclampsia-dependent genetic regulation at promoters"

**SUPPLEMENTARY MATERIAL**

**Supplementary Table 1. Quality control and sequencing metrics for samples in STRT Library 1.** Provided as a separate file.

**Supplementary Table 2. Quality control and sequencing metrics for samples in STRT Library 2.** Provided as a separate file.

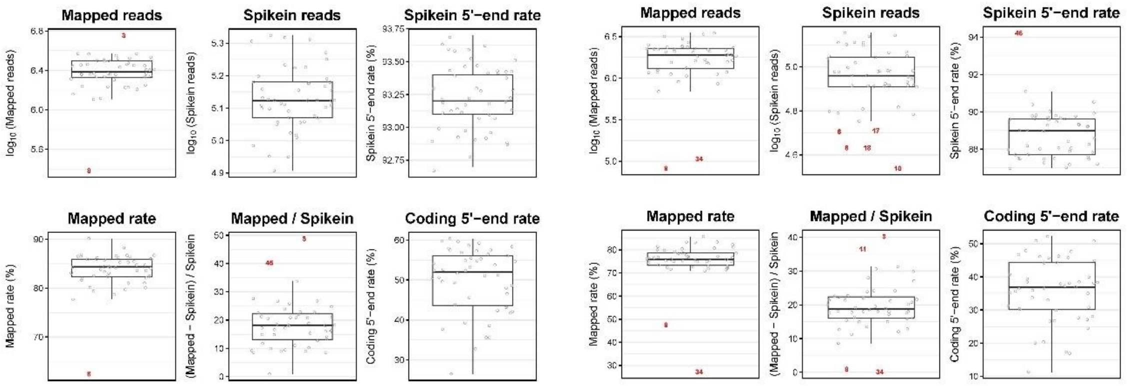


a) b)

**Supplementary Figure 1. Quality check plots for a) first STRT library, b) second STRT library.** In the first library, outliers were samples 3, 8, 45. In the second library, outliers were samples 3, 6, 8, 10, 11, 17, 18, 34 and 46.

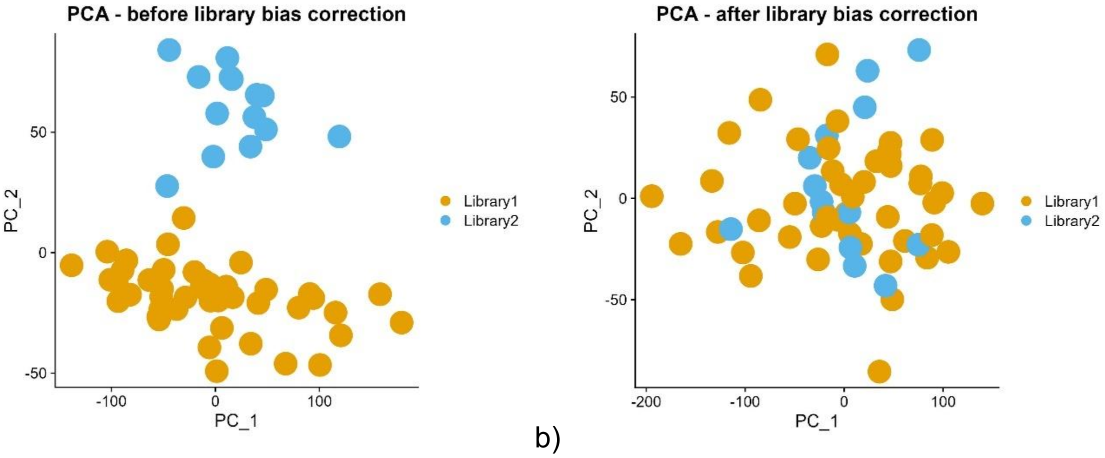


a) b)

**Supplementary Figure 2. PCA plots before and after library bias correction.**

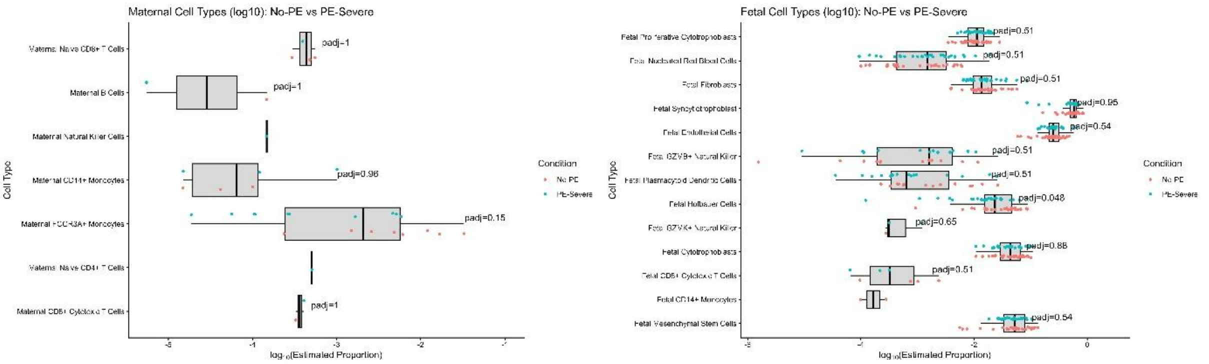


**Supplementary Figure 3. Estimated placental cell-type proportions are reduced in PE placental samples.** Estimated maternal and fetal cell-type proportions were compared between NP and PE groups. Proportions were log10-transformed for visualization. Differences were assessed using Wilcoxon rank-sum tests followed by Benjamini–Hochberg correction for multiple testing. Fetal Hofbauer cells showed a lower estimated proportion in PE placentas (padj=0.048), whereas no other cell type remained significant after multiple-testing correction.

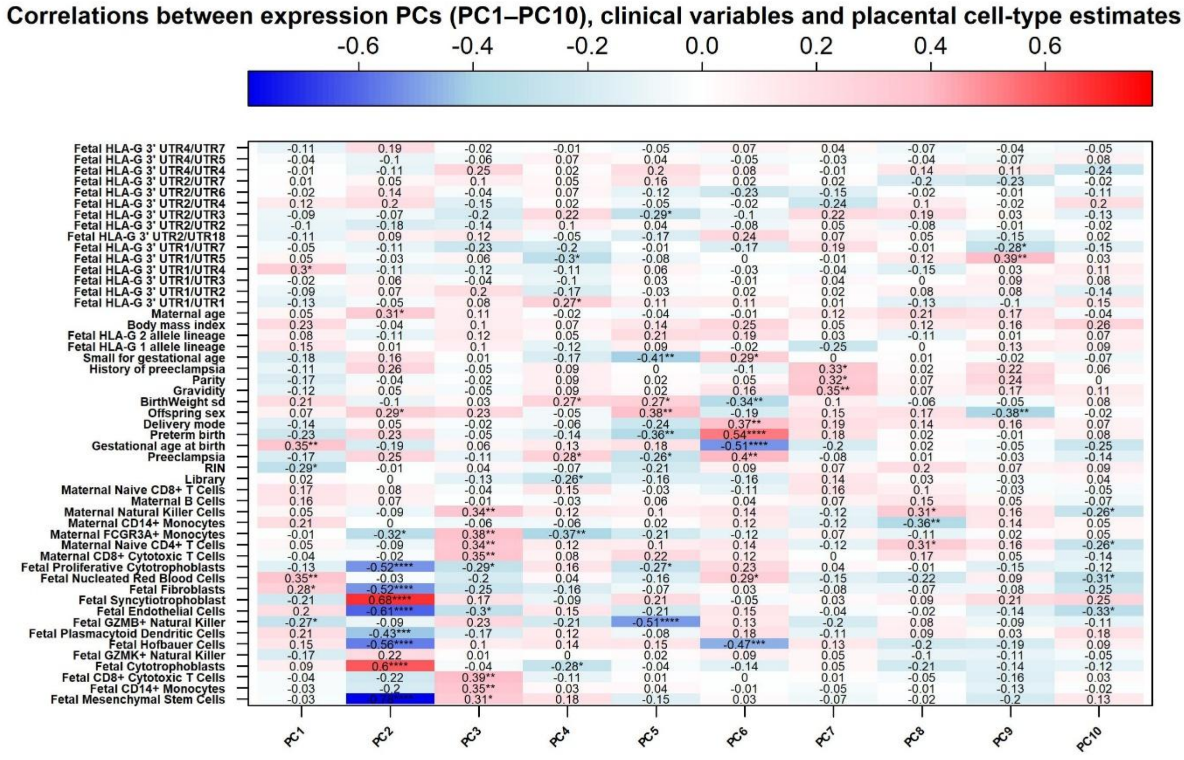


**Supplementary Figure 4. PE-associated gene-expression PCs also correlate with gestational and placental cell-type variables.** Heatmap showing associations between the first 10 principal components derived from gene-level expression profiles, clinical variables and estimated placental cell-type proportions. PE was associated with PC4 (r=0.28), PC5 (r=−0.26) and PC6 (r=0.40). Values show correlation coefficients. Asterisks indicate nominal significance: *p<0.05, **p<0.01, ***p<0.001, ****p<0.0001.

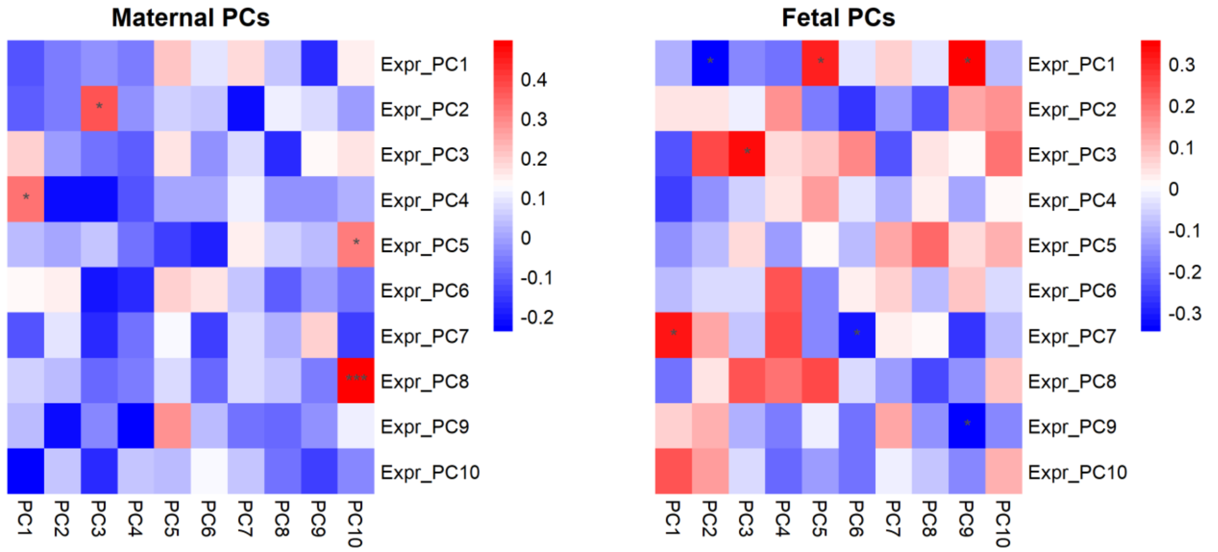


**Supplementary Figure 5. Correlations between genotype and gene-level expression PCs were calculated for eQTL covariate selection.** Heatmaps show Pearson correlations between the first 10 gene-level expression PCs and maternal (left) and fetal (right) genotype PCs. Expression PCs showing strong correlations with genotype PCs were excluded from the corresponding eQTL covariates to reduce collinearity. Asterisks indicate statistical significance: * p<0.05, ** p<0.01 and *** p<0.001.

**Supplementary Table 3. Significantly enriched immune related KEGG pathways among genes contributing to PE-associated expression PCs.** Only pathways with Benjamini-Hochberg adjusted p<0.05 were shown.

| **PCs** | **Eniched immune-related KEGG pathways** |
|---|---|
| PC4 | Oxidative phosphorylation, JAK-STAT singaling pathway, focal adhesion, phospatidylinositol signaling system |
| PC5 | Antigen processing and presentation, systemic lupus erythematosus, cell adhesion molecules CAMs |
| PC6 | Cell adhesion molecules CAMs, systemic lupus erythematosus, antigen processing and presentation, graft versus host disease, allograft rejection, type I diabetes mellitus, leukocyte transendothelial migration, natural killer cell mediated cytotoxicity, toll like receptor signaling pathway, oocyte meiosis |

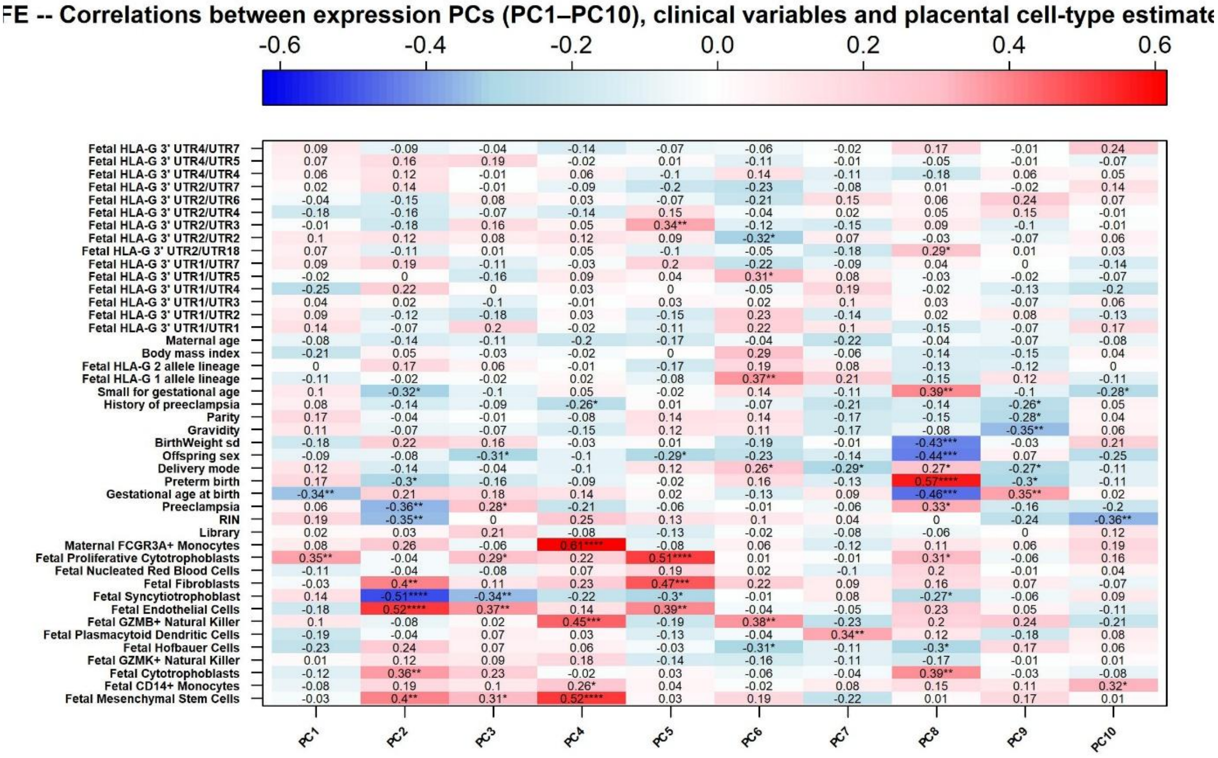


**Supplementary Figure 6. PE-associated promoter-expression PCs also correlate with clinical and placental cell-type variables.** Heatmap showing associations between the first 10 principal components derived from promoter-level expression profiles, clinical variables and estimated placental cell-type proportions. PE was associated with PC2 (r=−0.36), PC3 (r=0.28) and PC8 (r=0.33). Values show correlation coefficients. Asterisks indicate nominal significance: *p<0.05, **p<0.01, ***p<0.001, ****p<0.0001.

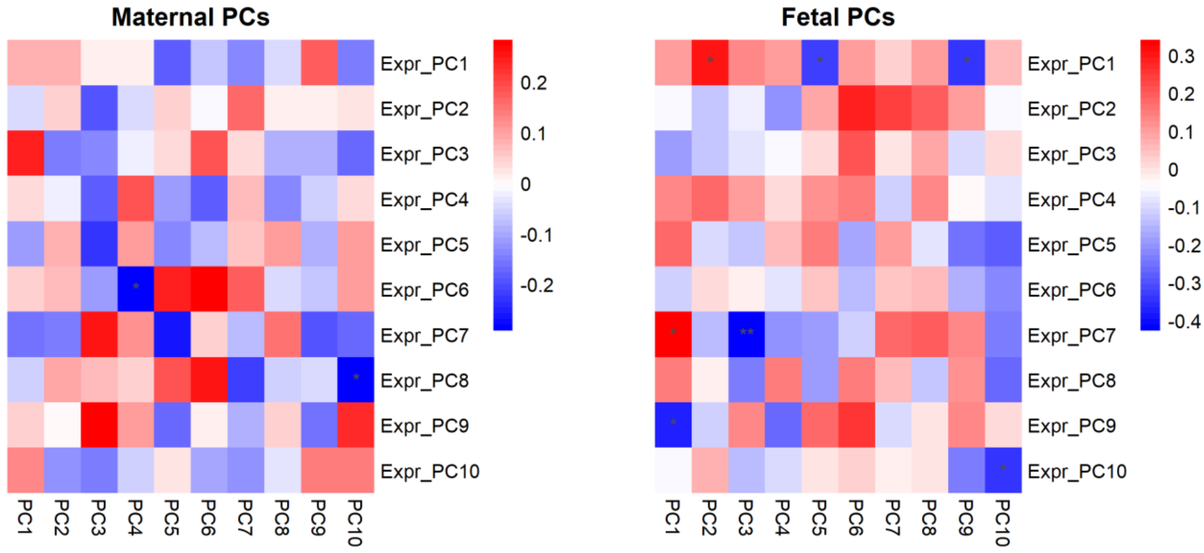


**Supplementary Figure 7. Genotype–TFE expression PC correlations informed covariate selection for maternal and fetal eQTL models.** Heatmaps show Pearson correlations between the first 10 promoter-level expression PCs and maternal (left) and fetal (right) genotype PCs. Expression PCs showing strong correlations with genotype PCs were excluded from the corresponding eQTL covariates to reduce collinearity. Asterisks indicate statistical significance: * p<0.05, ** p<0.01 and *** p<0.001.

**Supplementary Table 4. Top 20 suggestive maternal PE-dependent eQTL signals overlap regulatory regions.** The top 20 maternal gene-level interaction eQTL signals were ranked by genotype×PE interaction nominal pvalue. Candidate regulatory annotations were obtained from SCREEN and HaploReg and corresponding eQTL evidence in other tissues was assessed using GTEx.

| eGene | Gene description | Lead variant | Nearby cCREs (SCREEN) | Placental regulatory signals (Haploreg) | eQTL evidence in other tissues (GTEx) |
|---|---|---|---|---|---|
| *LILRB2* | leukocyte immunoglobulin like receptor B2 | rs3803902 | proximal enhancer | No | Yes |
| *CPLANE1* | ciliogenesis and planar polarity effector complex subunit 1 | rs3804449 | proximal enhancer, promoter | Yes, active enhancer region in placenta | Yes |
| *DOCK6* | dedicator of cytokinesis 6 | rs11666258 | distal enhancer | No | Yes |
| *TMCO1* | transmembrane and coiled-coil domains 1 | rs12027081 | TF, distal enhancer, CA-CTCF | No | No |
| *BNIP2* | BCL2 interacting protein 2 | rs66911717 | proximal enhancer, CA and TF | No | No |
| *XPNPEP1* | X-prolyl aminopeptidase 1 | Not available at dbSNP | | | |
| *CDH13* | cadherin 13 | rs8048006 | CA-CTCF, distal enhancer | No | No |
| *TDO2* | tryptophan 2,3-dioxygenase | rs13143482 | TF, distal enhancer | No | No |
| *NDUFAF7* | NADH:ubiquinone oxidoreductase complex assembly factor 7 | rs150305137 | Not available at the databases | | |
| *PI4K2A* | phosphatidylinositol 4-kinase type 2 alpha | rs11189444 | CA-CTCF, CA, distal enhancer, CA-TF | No | Yes |
| *ATP6V0E1* | ATPase H+ transporting V0 subunit e1 | rs11746431 | distal enhancer, distal enhancer | Yes, strong active enhancer region in placenta | Yes |
| *MSL3* | MSL complex subunit 3 | rs35992881 | CA | No | Yes |
| *TMEM50B* | transmembrane protein 50B | rs8130678 | distal enhancer | No | Yes |
| *ITM2C* | integral membrane protein 2C | rs1549573 | CA-CTCF, CA-CTCF, distal enhancer, CA-H3K4me3 | No | No |
| *CCNB1* | cyclin B1 | rs28451759 | distal enhancer, distal enhancer | No | Yes |
| *SPRED2* | sprouty related EVH1 domain containing 2 | rs1466649 | CA | No | No |
| *ECT2* | epithelial cell transforming 2 | rs67532538 | proximal enhancer, proximal enhancer, | Yes, transcription start site–associated | Not available |

| | | | | | |
|---|---|---|---|---|---|
| | | | promoter, distal enhancer | active promoter in placenta | |
| *AKR1E2* | aldo-keto reductase family 1 member E2 | rs7921950 | distal enhancer, proximal enhancer, TF | Yes, active enhancer region in placenta | Yes |
| *SEPTIN7* | septin 7 | rs34872540 | distal enhancer, CA | No | No |
| *SNRPA1* | small nuclear ribonucleoprotein polypeptide A' | rs3794504 | proximal enhancer, proximal enhancer, promoter | Yes, active promoter region (TSS-associated) in placenta | Yes |

Abbreviations: cCRE, candidate cis-regulatory element; CA, chromatin accessibility; CTCF, CCCTC-binding factor; TF, transcription factor; H3K4me1, histone H3 lysine 4 monomethylation; H3K4me3, histone H3 lysine 4 trimethylation; H3K27ac, histone H3 lysine 27 acetylation.

**Supplementary Table 5. Top 20 suggestive fetal PE-dependent eQTL signals overlap regulatory regions.** The top 20 fetal gene-level interaction eQTL signals were ranked by genotype×PE interaction nominal pvalue. Candidate regulatory annotations were obtained from SCREEN and HaploReg and corresponding eQTL evidence in other tissues was assessed using GTEx.

| **eGene** | **Gene description** | **Lead variant** | **Nearby cCREs (SCREEN)** | **Placental regulatory signals (Haploreg)** | **eQTL evidence in other tissues (GTEx)** |
|---|---|---|---|---|---|
| *LPCAT3* | lysophosphatidylcholine acyltransferase 3 | rs11056636 | CA-H3K4me3, proximal enhancer | No | No |
| *KHK* | ketohexokinase | rs35049337 | CA-CTCF, CA, CA-H3K4me3 | No | Not available |
| *CIAO1* | cytosolic iron-sulfur assembly component 1 | rs11681534 | distal enhancer | No | Yes |
| *NDUFAF5* | NADH:ubiquinone oxidoreductase complex assembly factor 5 | rs6105245 | distal enhancer and CA-H3K4m3 | No | Yes |
| *PHB2* | prohibitin 2 | rs7311336 | distal enhancer and TF | Yes, active enhancer region in placenta | Yes |
| *DPM1* | dolichyl-phosphate mannosyltransferase subunit 1, catalytic | rs6021247 | distal enhancer | No | No |
| *TNRC6C* | trinucleotide repeat containing adaptor 6C | rs117713991 | distal enhancer | No | No |
| *REXO2* | RNA exonuclease 2 | rs10790035 | distal enhancer and CA-H3K4m3 and CA-CTCF | No | No |
| *ANGEL2* | angel homolog 2 | rs4655315 | distal enhancer | No | Yes |

| | | | | | |
|---|---|---|---|---|---|
| *YJU2* | YJU2 splicing factor homolog | rs11667825 | CA, promoter, proximal enhancer | No | No |
| *COX5B* | cytochrome c oxidase subunit 5B | rs11681534 | distal enhancer | No | Yes |
| *CORO1C* | coronin 1C | rs1420664 | promoter, proximal enhancer, promoter | Yes, active promoter with enhancer-like activity in placenta!!! TssA chromatin state marked!!! | Yes |
| *TMED10* | transmembrane p24 trafficking protein 10 | rs61979196 | CA-H3K4me3 | No | Yes |
| *ESS2* | ess-2 splicing factor homolog | rs5993572 | proxomal enhancer, TF, proximal enhancer, distal enhancer, promoter | Yes, H3K4me3-marked region in placenta amnion | Yes |
| *PLAAT3* | phospholipase A and acyltransferase 3 | rs11231448 | no nearby cCREs found within 2000bp of it | No | Yes |
| *SLC35F6* | solute carrier family 35 member F6 | rs6711402 | CA, distal enhancer | No | Yes |
| *ANKRD39* | ankyrin repeat domain 39 | rs11681534 | distal enhancer | H3K27ac_Enh marked in placenta | Yes |
| *SMURF2* | SMAD specific E3 ubiquitin protein ligase 2 | Not available at dbSNP | | | |
| *CCNB1* | cyclin B1 | rs3730087 | proximal enhancer, promoter, promoter enhancer | Yes, H3K4me1_Enh | No |
| *ZFAND2A* | Zinc finger AN1-type containing 2A | rs9801230 | proximal enhancer, distal enhancer, proximal enhancer, CA-H3K4me3, promoter | Yes, H3K4me1_Enh marked in placenta (enhancer) | Yes |

Abbreviations: cCRE, candidate cis-regulatory element; CA, chromatin accessibility; CTCF, CCCTC-binding factor; TF, transcription factor; H3K4me1, histone H3 lysine 4 monomethylation; H3K4me3, histone H3 lysine 4 trimethylation; H3K27ac, histone H3 lysine 27 acetylation.

| Barcode | Qualified_reads | Total_reads | Redundancy | Mapped_reads | Mapped_rate | Spikein_reads | Spikein-5end_reads | Spikein-5end_rate | Coding_reads | Coding-5end_reads | Coding-5end_rate |
|---|---|---|---|---|---|---|---|---|---|---|---|
| STRT-PE1-1_1 | 2038231 | 1609141 | 1.26 | 1278784 | 79.4 | 80739 | 75638 | 93.6 | 820570 | 405153 | 49.3 |
| STRT-PE1-1_10 | 6060128 | 4361350 | 1.38 | 3698379 | 84.7 | 133079 | 124327 | 93.4 | 2817076 | 1193614 | 42.3 |
| STRT-PE1-1_11 | 4895730 | 3811671 | 1.28 | 3217662 | 84.4 | 129265 | 120910 | 93.5 | 2397530 | 943875 | 39.3 |
| STRT-PE1-1_12 | 4608401 | 3468428 | 1.32 | 2811799 | 81.0 | 147139 | 137203 | 93.2 | 1948294 | 840128 | 43.1 |
| STRT-PE1-1_13 | 4229737 | 3068337 | 1.37 | 2530649 | 82.4 | 151136 | 140904 | 93.2 | 1778522 | 744184 | 41.8 |
| STRT-PE1-1_14 | 2423235 | 1854383 | 1.30 | 1441012 | 77.7 | 151748 | 140692 | 92.7 | 910036 | 477709 | 52.4 |
| STRT-PE1-1_15 | 3442873 | 2557885 | 1.34 | 2014851 | 78.7 | 167464 | 155658 | 92.9 | 1391755 | 457014 | 32.8 |
| STRT-PE1-1_16 | 2944058 | 2214781 | 1.32 | 1775924 | 80.1 | 186657 | 174518 | 93.4 | 1036645 | 448772 | 43.2 |
| STRT-PE1-1_17 | 6286100 | 4095733 | 1.53 | 3550123 | 86.6 | 102152 | 95513 | 93.5 | 2919687 | 1600784 | 54.8 |
| STRT-PE1-1_18 | 4018706 | 2763215 | 1.45 | 2335770 | 84.5 | 123184 | 114897 | 93.2 | 1795927 | 819041 | 45.6 |
| STRT-PE1-1_19 | 5960221 | 3971005 | 1.50 | 3418254 | 86.0 | 132639 | 123619 | 93.1 | 2809047 | 1638047 | 58.3 |
| STRT-PE1-1_2 | 3069158 | 2070606 | 1.48 | 1707640 | 82.4 | 89037 | 83365 | 93.6 | 1206697 | 708119 | 58.6 |
| STRT-PE1-1_20 | 4906774 | 3325884 | 1.47 | 2885727 | 86.7 | 132763 | 124005 | 93.4 | 2320979 | 1310827 | 56.4 |
| STRT-PE1-1_21 | 5794493 | 3814472 | 1.51 | 3295330 | 86.3 | 142933 | 133161 | 93.1 | 2565186 | 1519265 | 59.2 |
| STRT-PE1-1_22 | 6130219 | 4289305 | 1.42 | 3711323 | 86.5 | 157835 | 146915 | 93.0 | 2882182 | 1460084 | 50.6 |
| STRT-PE1-1_23 | 4263948 | 3036216 | 1.40 | 2538958 | 83.6 | 178175 | 165701 | 92.9 | 1774036 | 827343 | 46.6 |
| STRT-PE1-1_24 | 3690688 | 2873582 | 1.28 | 2303537 | 80.1 | 210917 | 196401 | 93.1 | 1289704 | 343017 | 26.5 |
| STRT-PE1-1_25 | 5061414 | 3271188 | 1.54 | 2760163 | 84.3 | 96418 | 90026 | 93.3 | 2157909 | 1168249 | 54.1 |
| STRT-PE1-1_26 | 5670568 | 3661575 | 1.54 | 3159698 | 86.2 | 113936 | 105900 | 92.9 | 2336759 | 1392813 | 59.6 |
| STRT-PE1-1_27 | 3588994 | 2643489 | 1.35 | 2222789 | 84.0 | 128420 | 120016 | 93.4 | 1658339 | 806043 | 48.6 |
| STRT-PE1-1_28 | 3500017 | 2548043 | 1.37 | 2159596 | 84.7 | 126797 | 118449 | 93.4 | 1519279 | 730045 | 48.0 |
| STRT-PE1-1_29 | 2702785 | 2085464 | 1.29 | 1690392 | 81.0 | 150268 | 139989 | 93.1 | 979067 | 349336 | 35.6 |
| STRT-PE1-1_3 | 9424957 | 6448193 | 1.46 | 5688818 | 88.2 | 114206 | 106905 | 93.6 | 4404060 | 2247704 | 51.0 |
| STRT-PE1-1_30 | 4200918 | 2945279 | 1.42 | 2555930 | 86.7 | 132999 | 124458 | 93.5 | 1998193 | 1092668 | 54.6 |
| STRT-PE1-1_31 | 2764068 | 2110301 | 1.30 | 1682444 | 79.7 | 168336 | 156540 | 92.9 | 847098 | 301581 | 35.6 |
| STRT-PE1-1_32 | 5918991 | 3866028 | 1.53 | 3345941 | 86.5 | 204875 | 190696 | 93.0 | 2511362 | 1444102 | 57.5 |
| STRT-PE1-1_33 | 4077964 | 2699977 | 1.51 | 2275015 | 84.2 | 103916 | 96958 | 93.3 | 1697093 | 972769 | 57.3 |
| STRT-PE1-1_34 | 4519633 | 3178006 | 1.42 | 2711389 | 85.3 | 113685 | 106154 | 93.3 | 2174638 | 1202930 | 55.3 |
| STRT-PE1-1_35 | 3824419 | 2833988 | 1.34 | 2340172 | 82.5 | 132917 | 124212 | 93.4 | 1622057 | 711042 | 43.8 |
| STRT-PE1-1_36 | 3598194 | 2797753 | 1.28 | 2284795 | 81.6 | 143926 | 134118 | 93.1 | 1534921 | 579713 | 37.7 |
| STRT-PE1-1_37 | 5175416 | 3663562 | 1.41 | 3149590 | 85.9 | 142377 | 132901 | 93.3 | 2251781 | 949672 | 42.1 |
| STRT-PE1-1_38 | 4280393 | 2833807 | 1.51 | 2327248 | 82.1 | 155023 | 144082 | 92.9 | 1671623 | 936384 | 56.0 |
| STRT-PE1-1_39 | 4832588 | 3485603 | 1.38 | 2974621 | 85.3 | 149744 | 139695 | 93.2 | 2328394 | 900189 | 38.6 |
| STRT-PE1-1_4 | 2566849 | 1928490 | 1.33 | 1613163 | 83.6 | 117387 | 109711 | 93.4 | 1199977 | 615044 | 51.2 |
| STRT-PE1-1_40 | 3560379 | 2484336 | 1.43 | 2066127 | 83.1 | 202404 | 188071 | 92.9 | 1393352 | 705861 | 50.6 |
| STRT-PE1-1_41 | 3726386 | 2740141 | 1.35 | 2311364 | 84.3 | 105590 | 98783 | 93.5 | 1777364 | 985041 | 55.4 |
| STRT-PE1-1_42 | 4972647 | 3305965 | 1.50 | 2829737 | 85.5 | 119036 | 110910 | 93.1 | 2185985 | 1176975 | 53.8 |
| STRT-PE1-1_43 | 4898491 | 3361113 | 1.45 | 2887310 | 85.9 | 133916 | 125398 | 93.6 | 2209009 | 1278992 | 57.8 |
| STRT-PE1-1_44 | 3941439 | 2540009 | 1.55 | 2183553 | 85.9 | 112195 | 104554 | 93.1 | 1620173 | 836281 | 51.6 |
| STRT-PE1-1_45 | 6120361 | 4093777 | 1.49 | 3691121 | 90.1 | 89422 | 83602 | 93.4 | 3048871 | 1840361 | 60.3 |
| STRT-PE1-1_46 | 3900622 | 2641357 | 1.47 | 2294918 | 86.8 | 125654 | 117701 | 93.6 | 1833394 | 909475 | 49.6 |
| STRT-PE1-1_47 | 5594632 | 3642979 | 1.53 | 3127733 | 85.8 | 171039 | 159207 | 93.0 | 2256340 | 1183455 | 52.4 |
| STRT-PE1-1_48 | 3897431 | 2601585 | 1.49 | 2172051 | 83.4 | 207854 | 193914 | 93.2 | 1521174 | 836008 | 54.9 |
| STRT-PE1-1_5 | 2264766 | 1561696 | 1.45 | 1309327 | 83.8 | 130131 | 121258 | 93.1 | 791512 | 450585 | 56.9 |
| STRT-PE1-1_6 | 2778541 | 2107265 | 1.31 | 1729029 | 82.0 | 135566 | 126356 | 93.2 | 1124787 | 591987 | 52.6 |
| STRT-PE1-1_7 | 5129484 | 3306345 | 1.55 | 2856786 | 86.4 | 128489 | 119795 | 93.2 | 2305361 | 1376779 | 59.7 |
| STRT-PE1-1_8 | 470668 | 385764 | 1.22 | 241079 | 62.4 | 125340 | 117457 | 93.7 | 92614 | 53309 | 57.5 |
| STRT-PE1-1_9 | 5759633 | 3792195 | 1.51 | 3152277 | 83.1 | 117693 | 109295 | 92.8 | 1723175 | 917686 | 53.2 |

| Barcode | Qualified_reads | Total_reads | Redundancy | Mapped_reads | Mapped_rate | Spikein_reads | Spikein-5end_reads | Spikein-5end_rate | Coding_reads | Coding-5end_reads | Coding-5end_rate |
|---|---|---|---|---|---|---|---|---|---|---|---|
| STRT-PE1-2_1 | 2240071 | 1563609 | 1.43 | 1120605 | 71.6 | 91637 | 82369 | 89.8 | 801461 | 271138 | 33.8 |
| STRT-PE1-2_10 | 1965918 | 1336484 | 1.47 | 1115400 | 83.4 | 34539 | 30865 | 89.3 | 929360 | 257009 | 27.6 |
| STRT-PE1-2_11 | 4737032 | 2996871 | 1.58 | 2414279 | 80.5 | 64406 | 57345 | 89.0 | 1858277 | 379271 | 20.4 |
| STRT-PE1-2_12 | 4667696 | 3058227 | 1.52 | 2184592 | 71.4 | 117201 | 104647 | 89.2 | 1614246 | 437566 | 27.1 |
| STRT-PE1-2_13 | 4039062 | 2553796 | 1.58 | 1916237 | 75.0 | 88035 | 77356 | 87.8 | 1481646 | 415860 | 28.0 |
| STRT-PE1-2_14 | 2423575 | 1564338 | 1.54 | 1166007 | 74.5 | 62737 | 54583 | 87.0 | 890495 | 313609 | 35.2 |
| STRT-PE1-2_15 | 4686795 | 2826449 | 1.65 | 2053869 | 72.6 | 81646 | 71444 | 87.5 | 1706541 | 456608 | 26.7 |
| STRT-PE1-2_16 | 2424618 | 1648135 | 1.47 | 1184483 | 71.8 | 85316 | 74780 | 87.6 | 823208 | 202028 | 24.5 |
| STRT-PE1-2_17 | 3473765 | 1947042 | 1.78 | 1622621 | 83.3 | 51254 | 45882 | 89.5 | 1382009 | 613982 | 44.4 |
| STRT-PE1-2_18 | 2123574 | 1276839 | 1.66 | 1064163 | 83.3 | 42815 | 38429 | 89.7 | 898745 | 297441 | 33.0 |
| STRT-PE1-2_19 | 4199741 | 2282237 | 1.84 | 1878200 | 82.2 | 60836 | 54706 | 89.9 | 1631755 | 829838 | 50.8 |
| STRT-PE1-2_2 | 3118569 | 1697996 | 1.83 | 1322571 | 77.8 | 91246 | 81553 | 89.3 | 991796 | 517858 | 52.2 |
| STRT-PE1-2_20 | 3519205 | 2052807 | 1.71 | 1540987 | 75.0 | 96310 | 85212 | 88.4 | 1276879 | 584878 | 45.8 |
| STRT-PE1-2_21 | 4272250 | 2417614 | 1.76 | 1811163 | 74.9 | 95302 | 84029 | 88.1 | 1468066 | 680436 | 46.3 |
| STRT-PE1-2_22 | 4855314 | 2922881 | 1.66 | 2241728 | 76.6 | 95049 | 83388 | 87.7 | 1861240 | 665973 | 35.7 |
| STRT-PE1-2_23 | 3458870 | 2155016 | 1.60 | 1571967 | 72.9 | 82446 | 72148 | 87.5 | 1235224 | 381264 | 30.8 |
| STRT-PE1-2_24 | 4959451 | 3255921 | 1.52 | 2359286 | 72.4 | 99479 | 87278 | 87.7 | 1653452 | 187075 | 11.3 |
| STRT-PE1-2_25 | 6466407 | 3288517 | 1.96 | 2513665 | 76.4 | 112059 | 100440 | 89.6 | 2034510 | 829282 | 40.7 |
| STRT-PE1-2_26 | 5111469 | 2657402 | 1.92 | 2102783 | 79.1 | 104569 | 93927 | 89.8 | 1662238 | 844734 | 50.8 |
| STRT-PE1-2_27 | 3475882 | 2131680 | 1.63 | 1636923 | 76.7 | 96030 | 86979 | 90.5 | 1291014 | 492402 | 38.1 |
| STRT-PE1-2_28 | 4061617 | 2495629 | 1.62 | 1894157 | 75.8 | 112509 | 100448 | 89.2 | 1449538 | 507686 | 35.0 |
| STRT-PE1-2_29 | 3173513 | 2085033 | 1.52 | 1501158 | 71.9 | 82738 | 72732 | 87.9 | 1100559 | 186869 | 16.9 |
| STRT-PE1-2_3 | 6856456 | 3931226 | 1.74 | 3375438 | 85.8 | 81790 | 73342 | 89.6 | 2824974 | 1055768 | 37.3 |
| STRT-PE1-2_30 | 4003915 | 2331946 | 1.71 | 1835018 | 78.6 | 81162 | 70822 | 87.2 | 1431630 | 548369 | 38.3 |
| STRT-PE1-2_31 | 3377925 | 2352598 | 1.43 | 1668951 | 70.9 | 83916 | 73011 | 87.0 | 1093691 | 189338 | 17.3 |
| STRT-PE1-2_32 | 5158054 | 2803594 | 1.83 | 2113824 | 75.3 | 95490 | 84166 | 88.1 | 1753769 | 837543 | 47.7 |
| STRT-PE1-2_33 | 4939540 | 3006993 | 1.64 | 2274614 | 75.6 | 110738 | 99547 | 89.8 | 1860805 | 556470 | 29.9 |
| STRT-PE1-2_34 | 491023 | 391991 | 1.25 | 106650 | 27.2 | 95968 | 86102 | 89.7 | 8828 | 3350 | 37.9 |
| STRT-PE1-2_35 | 3208419 | 2032869 | 1.57 | 1543330 | 75.9 | 82246 | 74432 | 90.4 | 1104976 | 397504 | 35.9 |
| STRT-PE1-2_36 | 4994520 | 2877800 | 1.73 | 2211090 | 76.8 | 94913 | 84478 | 89.0 | 1732568 | 526943 | 30.4 |
| STRT-PE1-2_37 | 5930934 | 2855693 | 2.07 | 2319747 | 81.2 | 82681 | 73337 | 88.6 | 1987729 | 893828 | 44.9 |
| STRT-PE1-2_38 | 4581422 | 2644479 | 1.73 | 1910594 | 72.2 | 90130 | 79590 | 88.3 | 1565807 | 627511 | 40.0 |
| STRT-PE1-2_39 | 5431823 | 3014694 | 1.80 | 2202717 | 73.0 | 116196 | 101419 | 87.2 | 1784530 | 681353 | 38.1 |
| STRT-PE1-2_4 | 1717645 | 1110741 | 1.54 | 857557 | 77.2 | 65665 | 58669 | 89.3 | 680297 | 262559 | 38.5 |
| STRT-PE1-2_40 | 4832974 | 2994055 | 1.61 | 2280130 | 76.1 | 108398 | 95399 | 88.0 | 1794914 | 561845 | 31.3 |
| STRT-PE1-2_41 | 5631378 | 3499357 | 1.60 | 2699967 | 77.1 | 138064 | 124327 | 90.0 | 2137898 | 569979 | 26.6 |
| STRT-PE1-2_42 | 7561765 | 4329899 | 1.74 | 3502852 | 80.8 | 139219 | 125154 | 89.8 | 2942931 | 1089146 | 37.0 |
| STRT-PE1-2_43 | 7932793 | 3996240 | 1.98 | 3196728 | 79.9 | 142830 | 130171 | 91.1 | 2633530 | 1261537 | 47.9 |
| STRT-PE1-2_44 | 4588066 | 2986224 | 1.53 | 2240322 | 75.0 | 129484 | 116031 | 89.6 | 1717512 | 344956 | 20.0 |
| STRT-PE1-2_45 | 4804912 | 2753976 | 1.74 | 2071287 | 75.2 | 126405 | 112617 | 89.0 | 1603894 | 617183 | 38.4 |
| STRT-PE1-2_46 | 2482043 | 1523679 | 1.62 | 1240890 | 81.4 | 129095 | 121659 | 94.2 | 822582 | 300063 | 36.4 |
| STRT-PE1-2_47 | 7537315 | 3782307 | 1.99 | 2973571 | 78.6 | 129032 | 112920 | 87.5 | 2499314 | 1105889 | 44.2 |
| STRT-PE1-2_48 | 5507246 | 3176096 | 1.73 | 2388615 | 75.2 | 125692 | 109805 | 87.3 | 1870196 | 686077 | 36.6 |
| STRT-PE1-2_5 | 2010133 | 1152806 | 1.74 | 904591 | 78.4 | 56624 | 49864 | 88.0 | 731717 | 337658 | 46.1 |
| STRT-PE1-2_6 | 1344464 | 944291 | 1.42 | 693247 | 73.4 | 50696 | 44368 | 87.5 | 497159 | 150343 | 30.2 |
| STRT-PE1-2_7 | 2618998 | 1475489 | 1.77 | 1180493 | 80.0 | 61648 | 53989 | 87.5 | 991725 | 463372 | 46.7 |
| STRT-PE1-2_8 | 224212 | 171746 | 1.30 | 81905 | 47.6 | 42873 | 37816 | 88.2 | 34068 | 13438 | 39.4 |
| STRT-PE1-2_9 | 5826407 | 3086393 | 1.88 | 2378612 | 77.0 | 81992 | 73518 | 89.6 | 1873386 | 857046 | 45.7 |
